# Dynamically unstable ESS in matrix games under time constraints

**DOI:** 10.1101/2021.08.05.455237

**Authors:** Tamás Varga, József Garay

## Abstract

Matrix games under time constraints are natural extensions of matrix games. They consider the fact that, in addition to the payoff, a pairwise interaction has a further consequence for the contestants. Namely, both players have to wait for some time before becoming fit to participate in a subsequent interaction. Every matrix game can be assigned a continuous dynamical system (the replicator equation) which describes how the frequencies of different phenotypes evolve in the population. One of the fundamental theorems of evolutionary matrix games asserts that the state corresponding to an evolutionarily stable strategy is an asymptotically stable rest point of the replicator equation (Taylor and Yonker 1978, Hofbauer et al. 1979, Zeeman 1980). Garay et al. (2018) and Varga et al. (2020) generalized the statement to two-strategy and, in some particular cases, three- or more strategy matrix games under time constraints. However, the question of whether the implication holds in general remained open. Here examples are provided demonstrating that the answer is no. Moreover, we point out through the rock-scissor-paper game that arbitrary small differences between waiting times can destabilize the rest point corresponding to an ESS. It is also shown that a stable limit cycle can arise around the unstable rest point in a supercritical Hopf bifurcation.

**Mathematics Subject Classification:** 91A22, 92D15, 92D25, 91A80, 91A05, 91A10, 91A40, 92D40

## 1 Introduction

Classic evolutionary matrix games introduced by Maynard Smith and Price (1973) describe populations where individuals compete with each other through pairwise interactions (Maynard Smith 1982, Chapter 6-7 in Hofbauer and Sigmund 1998, Chapter 2.3 in Mesterton-Gibbons 2001, Chapter 6 in Broom and Rychtář 2013). As a result of an interaction, the contestants get some payoff depending on their strategies/phenotypes. The payoffs of the consecutive interactions then determine the (relative) fitness, thereby the evolutionary success, of a given phenotype. However, in a lot of situations the consequence of an interaction is not only some payoff but some time the contestants have to wait so as to become fit for a subsequent interaction (e.g. to recover from an injury, to handle the payoff (food), to collect energy after a long fight, etc.). Therefore a part of the population is unable to interact in a given moment and there is no intake during waiting time. These can significantly influence the fitness and so the evolutionary outcomes (Garay et al. 2017, Křivan and Cressman 2017). For example, in the classical Prisoner’s dilemma, the ESS strategy is the defector strategy but if the waiting time related to the interaction of defectors is long enough then the average intake of defectors is smaller than that of cooperators resulting in the evolutionary stability of the cooperator strategy in that model. Similarly, if the waiting time after a Hawk-Hawk interaction in the Hawk-Dove game is long enough then pure Dove strategy can be ESS even at a low cost of fighting. These examples well demonstrate that the consideration of time constraints in game theoretical models can lead to surprising phenomena suggesting the necessity of further mathematical investigations.

Maynard Smith and Price (1973) studied monomorphic populations where every individual initially has the same (pure or mixed) phenotype. For this case, Maynard Smith introduced the following verbal static definition: “An ESS is a strategy such that, if all the members of a population adopt it, then no mutant strategy could invade the population under the influence of natural selection” (Maynard Smith 1982, p. 10). This condition means that the ESS is always fixed by the natural selection if the frequency of mutants is small.

Later Taylor and Yonker (1978) suggested a polymorphic approach when a population is considered that consists of individuals of *N* different genetically fixed pure phenotypes. Then the temporal evolution of the population is modelled by some dynamics, often the replicator equation (Taylor and Yonker 1978, Hofbauer and Sigmund 1998), and a locally asymptotically stable rest point of the dynamics is the endpoint of the evolution.

It is a central result of classic evolutionary game theory that, in the case of matrix games, the corresponding state of an ESS is a locally asymptotically stable rest point of the replicator equation (Taylor and Yonker 1978, Hofbauer et al. 1979, Zeeman 1980). The relationship holds not only for the matrix games but for some cases when the fitness function is not bilinear in the phenotype of the focal individual and the population average strategy. Indeed, it is enough to assume the linearity in the strategy of the focal individual (see e.g. Thomas 1985, Chapter 7.2 in Hofbauer and Sigmund 1998 or Cressman 2003). For other fitness functions, however, the validity of an analogous theorem is an open question.

In the case of evolutionary matrix games under time constraints, the fitness function is neither linear in the strategy of the focal individual nor in the average strategy of the population. Similarly to the aforementioned classic result, Garay et al. (2018) succeeded in proving the asymptotic stability of the rest point corresponding to an ESS for two-strategy games (Corollary 4.3 in Garay et al. 2018). The proof exploit that the set of the convex combinations of the strategies for which the replicator equation is considered is one-dimensional. The higher dimensional case, however, remained open except some special cases (Theorem 4.1 in Garay et al. 2018, Theorem 4.7 and Theorem 4.9 in Varga et al. 2020) when the corresponding state is asymptotically stable or at least stable. Nevertheless, the authors’ conjecture was that the stability does not hold in general.

In this paper we show that the conjecture is true giving examples for matrix games under time constraints with evolutionary stable strategy such that the corresponding state of the replicator equation is unstable even for 3 dimensional strategies.

One of our example starts from the rock-scissor-paper game. This game models cyclic dominance (Sinervo and Lively 1996, Kerr et al. 2002, Liao et al. 2019) which can result in the cyclic coexistence of three phenotypes (Hofbauer and Sigmund 1998, Nowak 2006, Mobilia 2010, Toupo and Strogatz 2015). We show that arbitrary small distinctions between waiting times can destabilize the corresponding state. Moreover, we point out that a limit cycle can arise around it in a Hopf bifurcation which is again a new phenomenon compared to classic matrix games (Zeeman 1980, Bomze 1983).

## 2 Preliminaries

We introduce some functions and notations necessary to our calculations. For the detailed backgrounds (the heuristic thoughts and the mathematical derivation of the formulas), we refer the reader to Garay et al. (2017) and Garay et al. (2018). Let *S*_*N*_ denote the set

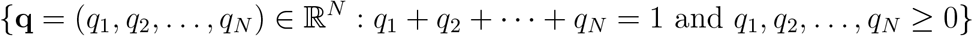

(*N* is a positive integer). The elements of *S*_*N*_ are called strategies or (pheno)types. The special strategies **e**_1_ = (1, 0, …, 0), **e**_2_ = (0, 1, 0, …, 0), …, **e**_*N*−1_ = (0, 0, …, 0, 1, 0), **e**_*N*_ = (0, 0, …, 0, 1) are called the pure strategies/(pheno)types. Obviously, every strategy **q** is a convex combination or, in game theoretical terminology, the mix of the pure strategies. Sometimes, instead of saying “individual following strategy **q**”, we just say **q** individual or **q**-strategist.

We consider a huge well mixed population of **q**_1_, **q**_2_, …, **q**_*n*_ individuals (mathematically the number of individuals tends to infinity). The individuals look for opponents to make a pairwise interactions. An interaction implies two consequences for the participants: some **payoff** and some **waiting time** which is to wait before the individual becomes suitable for a next interaction. This waiting time can correspond to the handling time in optimal foraging theory (the time necessary for processing a pray item, Charnov 1976, Garay et al. 2012), the recovery time from an injury acquired in the interaction, the time necessary to collect energy to a newer interaction, etc. During waiting time the individual is unable to interact, therefore it is said to be **inactive** while an individual searching for an opponent is called **active**. If an individual uses the *i*-th pure strategy and the opponent uses the *j*-th pure strategy in an interaction then the payoff of the focal individual is *a*_*ij*_. The *N* × *N* matrix *A* = (*a*_*ij*_) is called the **payoff matrix**. Similarly, *τ*_*ij*_ denotes the waiting time the focal individual should wait after such an interaction and the *N* × *N* matrix *T* = (*τ*_*ij*_) is called the **time-constraint matrix**. It is assumed that *τ*_*ij*_ ≥ 0. Both the searching time and the waiting time is an exponential random variable. The expected value of the searching time is set to be 1 (after normalization) while the expected value of the waiting time is *τ*_*ij*_ depending on the strategies of the opponents as mentioned before. (The interaction itself is considered to be instantaneous or it can be considered the part of the waiting time.)

If the searching time ends, the searching individual randomly chooses an opponent. If the opponent is another searching individual then interaction occurs, if the opponent is just waiting, then nothing occurs but the searching time of the searching individual starts over. With these assumptions, a Markov model can be built which has a unique stationary distribution (Garay et al. 2017). If first the time of observation then the number of individuals tends to the infinity such that the frequency of the **q**_*i*_ individuals tends to *x*_*i*_ and *W*_*i*_(**x**) denotes the limit of the average intake of a **q**_*i*_ individual per time unit with **x** = (*x*_1_, *x*_2_, …, *x*_*n*_) then

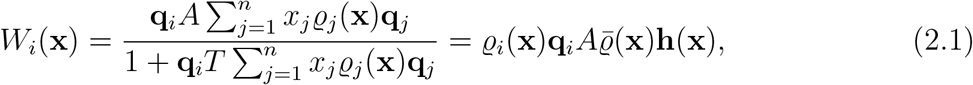

where *ϱ*_*i*_(**x**) is the unique solution in [0, 1] to the equation system:

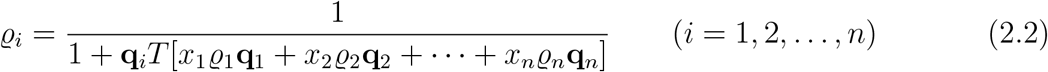

(Theorem 1 and Lemma 2 in Garay et al. 2017) and

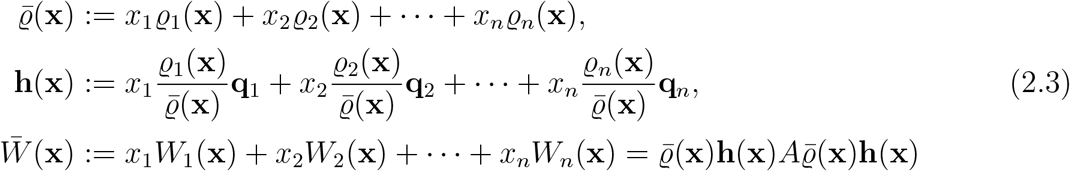

(If a strategy **q** ∈ *S*_*N*_ is multiplied by an *N* × *N* matrix M from the right then **q** should be considered a row vector while if it is multiplied by M from the left then **q** should be considered a column vector.) *W*_*i*_(**x**) represents the fitness of a **q**_*i*_ individual in the population. The numerator is just the expected payoff of an interaction (including the case when the opponent is in waiting time so there is no real interaction). The denominator is the expected length of the time between two consecutive interactions: it consists of the time necessary for searching and the waiting time. *W*_*i*_ can thus be viewed as an intake rate in essence. To see a possible intuitive interpretation of *ϱ*_*i*_ one should recall that 1 is corresponds to the average duration of a searching period, so *ϱ*_*i*_ gives the proportion of the life time of a **q**_*i*_ individual which is spent searching for an opponent, in other words, in active state. There is another meaning. It gives the proportion of active individuals in the subpopulation of **q**_*i*_ individuals in stationary state if the population size tends to infinity. [This is ensured by the fact that the underlying Markov model is ergodic (Theorem 1 in Garay et al. 2017).] Consequently, 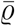 is the proportion of active individuals in the whole population while **h**(**x**) is the average strategy of an individual in the subpopulation of active individuals.

In the previous paragraphs, we have reviewed the expressions related to the polymorphic case. Now we turn to the monomorphic one. If **q** ∈ *S*_*N*_ is a strategy then *ρ*(**q**) denotes the unique solution in [0, 1] to the equation

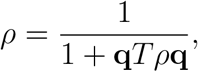

that is,

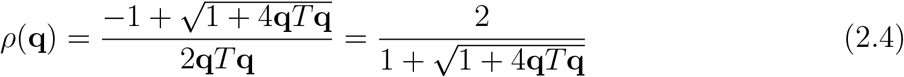

(Lemma 2 in Garay et al. 2017). Then

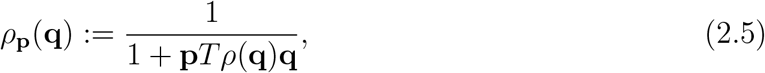

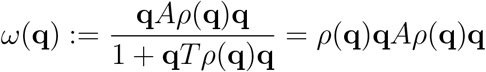 and 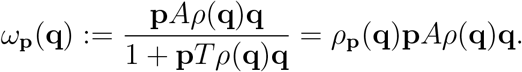 It follows that both *ρ*(**q**) and *ρ*_**p**_(**q**) has continuous second order partial derivatives with respect to the coordinates of **q** = (*q*_1_, *q*_2_, …, *q*_*N*_) ∈ *S*_*N*_. Analogously to the polymorphic case, *ρ*(**q**) can be interpreted as the proportion of the active individuals in a population of individuals following strategy **q**. *ω*(**q**) represents the fitness of an individual in this population while *ω*_**p**_(**q**) can be considered as the fitness of a single **p** individual in a large population of **q** individuals (as if a single **p**-strategist were in an infinitely large population of **q** individuals where *ρ*_**p**_ is the probability that the **p** individual is active in a given moment).

If **p, q** ∈ *S*_*N*_ and *ε* ∈ [0, 1] let *ρ*_**p**_(**p, q**, *ε*) and *ρ*_**q**_(**p, q**, *ε*), respectively, be the unique solution in [0, 1] to the equation system

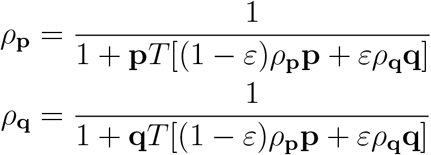

(Lemma 2 in Garay et al. 2017). *ρ*_**p**_ and *ρ*_**q**_, respectively, are the proportion of active individuals in the subpopulation of phenotype **p** and phenotype **q**, respectively, in a population where (1 − *ε*) is the frequency of **p** individuals and *ε* is that of **q** individuals. The fitness of these types is described as follows

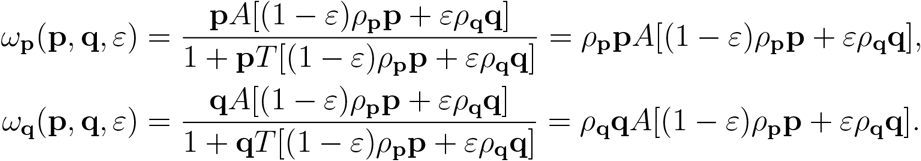

Note that *ρ*(**q**) = *ρ*_**q**_(**q, p**, 0) = *ρ*_**q**_(**p, q**, 1), *ω*(**q**) = *ω*_**q**_(**q**) = *ω*_**q**_(**q, p**, 0) = *ω*_**q**_(**p, q**, 1), *ρ*_**p**_(**q**) = *ρ*_**p**_(**p, q**, 1) = *ρ*_**p**_(**q, p**, 0) and *ω*_**p**_(**q**) = *ω*_**p**_(**p, q**, 1) = *ω*_**p**_(**q, p**, 0) for any **p, q** ∈ *S*_*N*_ and *ε* ∈ [0, 1].

We quote some statements crucial in understanding the relationship between the monomorphic and the polymorphic model.

### Lemma 2.1 (Garay et al. 2018, Proposition 3.1)

*Consider a polymorphic population of phenotypes* **q**, **q**, …, **q** *with frequency distribution* (*x, x*, …, *x*). *Then* 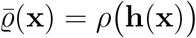 *and*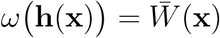, *that is the fitness of an* **h**(**x**) *individual is just the average fitness* 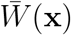 *of the phenotypes* **q**_1_, **q**_2_, …, **q**_*n*_ *in the polymorphic model*.

It is clear for classical matrix games that if **p** = (*p*_1_, *p*_2_, …, *p*_*N*_) is a strategy then (*x*_1_, *x*_2_, …, *x*_*N*_) with *x*_*i*_ = *p*_*i*_ is the corresponding state in the polymorphic systems of pure phenotypes **e**_1_, **e**_2_, …, **e**_*N*_ in the sense that ∑_*i*_ *x*_*i*_**e**_*i*_ = **p**. Consequently, if **p** is an ESS then it is natural to investigate the stability property of the state (*x*_1_, *x*_2_, …, *x*_*N*_) with *x*_*i*_ = *p*_*i*_ for the replicator equation with respect to **e**_1_, **e**_2_, …, **e**_*N*_. If the replicator equation is considered for arbitrary strategies **q**_1_, **q**_2_, …, **q**_*n*_ then it is also clear that the corresponding state(s) of **p** is (are) the state(s) (*x*_1_, *x*_2_, …, *x*_*n*_) with **p** = *x*_1_**q**_1_ + *x*_2_**q**_2_ + ⋯ + *x*_*n*_**q**_*n*_, that is, the state(s) at which the average strategy of the population is **p**. For matrix games under time constraints, however, the situation is generally not so obvious. The answer can be found in the next lemma of Garay et al. (2018) which asserts that the corresponding state is the state at which the average strategy of active individuals is **p**.

### Lemma 2.2 (Garay et al. 2018, Corollary 6.3)

*Consider a population of phenotypes* **q**_1_, **q**_2_,…, **q**_*n*_ ∈ *S*_*N*_. *Assume that* **r** = *θ*_1_**q**_1_ +*θ*_2_**q**_2_ +…+*θ*_*n*_**q**_*n*_ *with some* ***θ*** = (*θ*_1_, *θ*_2_, …, *θ*_*n*_) ∈ *S*_*n*_.

*If*

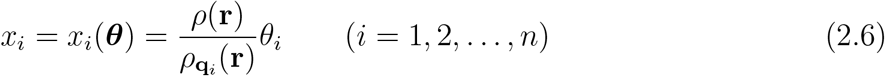

*then* **h**(**x**) = **r** *where* **x** = **x**(***θ***) = *x*_1_(***θ***), *x*_2_(***θ***), …, *x*_*n*_(***θ***), *that is, the average strategy of active individuals in a population of* **q**_1_, **q**_2_,…, **q**_*n*_ *individuals with frequencies x*_1_(***θ***), *x*_2_(***θ***),…, *x*_*n*_(***θ***) *is θ*_1_**q**_1_ + *θ*_2_**q**_2_ + ⋯ + *θ*_*n*_**q**_*n*_ = **r**. *Moreover*, **x**(***θ***) *is the unique state for which* 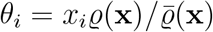 (*i* = 1, 2, …, *n*) *holds*.

We consider two particular cases. One of them is when *n* = *N* = 3, **q**_1_ = **e**_1_, **q**_2_ = **e**_2_ and **q**_3_ = **e**_3_. Let **r** = (*r*_1_, *r*_2_, 1 − *r*_1_ − *r*_2_) = *r*_1_**e**_1_ + *r*_2_**e**_2_ + (1 − *r*_1_ − *r*_2_)**e**_3_ be a strategy from *S*_3_. By the previous lemma, there exits a unique^1^ state **x**(**r**) ∈ *S*_3_ with **h x**(**r**) = **r**, namely, 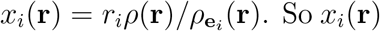 has continuous partial derivatives with respect to the variables *r*_1_ and *r*_2_. The notation **x**^*R*^(**r**) is used (“R” refers to “restricted”) if necessary to emphasize that **x**(**r**) is concerned with this case.

The other case is when *N* = 3, *n* = 4, **q**_1_ = **e**_1_, **q**_2_ = **e**_2_, **q**_3_ = **e**_3_ and **q**_4_ = **p**\* := (1/3, 1/3, 1/3). Then most of the strategies from *S*_3_, namely, the strategies in the interior of *S*_3_ can be expressed in the form *θ*_1_**e**_1_ + *θ*_2_**e**_2_ + *θ*_3_**e**_3_ + *θ*_4_**p**\* in more than one way. For such a strategy **r**, there are thus more then one state **x** = (*x*_1_, *x*_2_, *x*_3_, *x*_4_) with **h**(**x**) = **r**. So it would be ambiguous the notation **x**(**r**) for a state with **h**(**x**(**r**)) = **r**. Therefore we associate the coefficient-list ***θ*** = (*θ*_1_, *θ*_2_, *θ*_3_, *θ*_4_) from the convex combination *θ*_1_**e**_1_ + *θ*_2_**e**_2_ + *θ*_3_**e**_3_ + *θ*_4_**p**\* = **r** to the state **x** = **x**(***θ***) = **x**(*θ*_1_, *θ*_2_, *θ*_3_, *θ*_4_) with 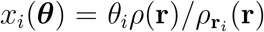 rather than associating a strategy **r** to such a state. If it is necessary to emphasize that **x**(***θ***) is concerned with this case then we use the notation **x**^*E*^(***θ***) (“E” refers to “extended”). Observe that if the vector (*x*_1_, *x*_2_, 1 − *x*_1_ − *x*_3_, 0) ∈ *S*_4_ is identified with the vector (*x*_1_, *x*_2_, 1 − *x*_1_ − *x*_3_) ∈ *S*_3_, then the map **x**^*R*^ is just the restriction of the map **x**^*E*^ to the set *S*_3_.

Note the following “exchange rule” between strategies and states. This is just the simple consequence of the preceding lemmas and the definitions of the expressions above.

### Corollary 2.3 (Exchange rule)

*Let* **r** *be a strategy in S*_3_ *and consider the polymorphic population of the pure phenotypes* **e**_1_, **e**_2_, **e**_3_. *Then the following relationships hold:*

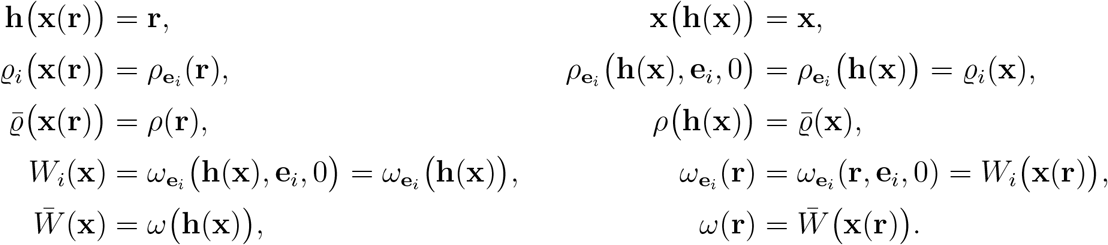

Using the notations above the evolutionary stability of a strategy is defined as follows.

### Definition 2.4

*A strategy* **p**\* *is uniformly evolutionary stable strategy of the matrix game under time constraints (UESS in short) if there is an ε*_0_ > 0 *such that*

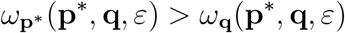

*for any strategy* **q** ≠ **p**\* *and* 0 < *ε* < *ε*_0_.

The adverb “uniformly” is used because *ε*_0_ does not depend on **q**. If *ε*_0_ can depend on **q** then **p**\* is called ESS without the adverb “uniformly”. Clearly, if **p**\* is a UESS then **p**\* is an ESS too. The converse is true in the case of classic matrix games (without time constraints) (Theorem 6.4.1 in Hofbauer and Sigmund 1998) but it is an open question for matrix games under time constraints. We do not want to deal with this problem in this article only we wanted to explain the cause of using the term “UESS”. This distinction between UESS and ESS naturally emerges for non-linear fitness functions (Bomze and Weibull 1995) and UESS is more appropriate for those (Pohley and Thomas 1983) to describe evolutionary stability.

We look for examples having a UESS **p**\* such that the corresponding state **x**(**p**\*) is unstable rest point of the replicator equation with respect to the pure strategies.

## 3 Dynamically unstable UESS

The concept of UESS described in Definition 2.4 mathematically tries to express the intuitive expectation that an evolutionary stable strategy must be proof against invasion. This static definition is very useful tool in applied evolutionary game theory because it permits us to predict the evolutionary outcome without analyzing a sophisticated model which describes the temporal changes in a population. However, from the point of view of theoretical evolutionary game theory, we need some dynamics describing the temporal evolution of the population (cf. Introduction in Cressman 1992). The usual dynamics is the replicator equation introduced by Taylor and Yonker (1978) in evolutionary game theory. If the population consists of **q**_1_, **q**_2_, …, **q**_*n*_ individuals with frequencies *x*_1_, *x*_2_, …, *x*_*n*_ then the corresponding replicator equation is the following:

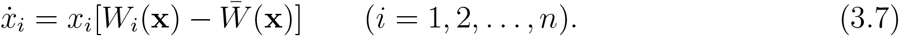

The question arises: what is the connection between the UESS concept and the dynamical stability. It is well known in the case of classic evolutionary matrix games^2^ that an UESS always implies the asymptotic stability of the corresponding equilibrium point of the associated replicator equation (Taylor and Yonker 1978, Hofbauer et al. 1979, Zeeman 1980 or Theorem 7.2.4 in Hofbauer and Sigmund 1998).

An analogues implication for matrix games under time constraints up to now was just proved when the the strategy space consists of 2 dimensional strategies. For higher dimensional strategies, however, the question remained open in general (Garay et al. 2018, Varga et al. 2020).

Now we give examples exhibiting UESS with unstable corresponding equilibrium, that is the accustomed implication of the classical model does not hold in the more general model of matrix games under time constraints. This shows that the consideration of time effects can dramatically alter the outcome of evolution.

The strategy space of the examples is

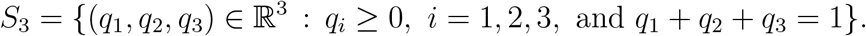

In each example, strategy **p**\* = (1/3, 1/3, 1/3) is a UESS which is verified i n section A.1 and A.2. The corresponding state is **x** = **x**(**p**\*) = **x**^*R*^(**p**\*) = (*x*_1_(**p**\*), *x*_2_(**p**\*), *x*_3_(**p**\*)) for which **h**(**x**) = **p**\* (see Lemma 2.2) (recall the notations **x**^*R*^ and **x**^*E*^ in the second paragraph after Lemma 2.2). We analyse the replicator equation (3.7) for the polymorphic population consisting of pure phenotypes **e**_1_, **e**_2_, **e**_3_ with frequencies *x*_1_, *x*_2_ and *x*_3_ = 1 − *x*_1_ − *x*_2_:

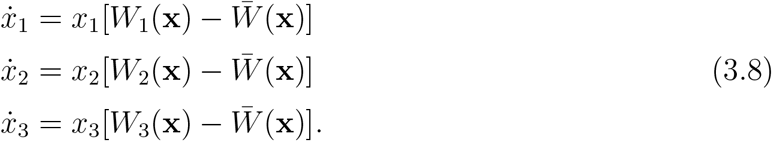

We also investigate the replicator equation with respect to the population of **q**_1_ = **e**_1_, **q**_2_ = **e**_2_, **q**_3_ = **e**_3_ and **q**_4_ = **p**\* = (1/3, 1/3, 1/3) individuals. This means the extension of the previous dynamics (3.8) adding a forth equation:

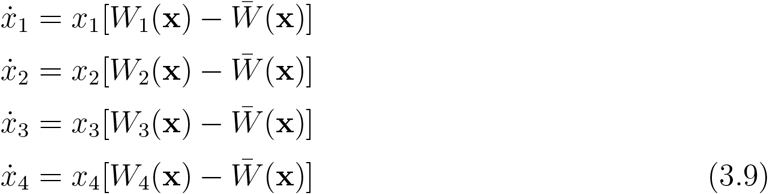

where *x*_4_ describes the frequency of **p**\* individuals. The phase space of this dynamics is *S*_4_. State (1, 0, 0, 0) corresponds to pure strategy **e**_1_ in the sense that in this state every individual follows strategy **e**_1_. Similarly, state (0, 1, 0, 0) corresponds to pure strategy **e**_2_, state (0, 0, 1, 0) corresponds to pure strategy **e**_3_ and state (0, 0, 0, 1) corresponds to strategy **p**\*. Clearly, if the replicator equation (3.9) is restricted to the face with vertices (1, 0, 0, 0), (0, 1, 0, 0) and (0, 0, 1, 0) we get back the dynamics (3.8). Accordingly, this face contains the state **x**^*E*^(1/3, 1/3, 1/3, 0) (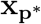 for short) corresponding to the UESS **p**\* in which each individual follows one of the three pure strategies and the average strategy of active individuals is the UESS **p**\*. Note that this state is the same as the state **x**^*R*^(**p**\*) for the replicator equation with respect to pure strategies **e**_1_, **e**_2_, **e**_3_.

By Lemma 2.2, the segment (the green segment in Figure 2 and 5) connecting 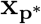 with (0, 0, 0, 1) precisely consists of the states corresponding to the UESS **p**\* and the coordinates of these states are

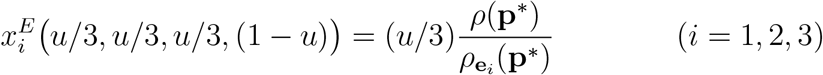

and

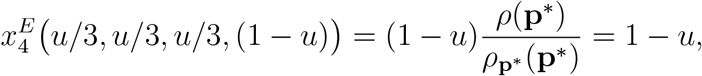

respectively, where *u* runs over the interval [0, 1]. These states are equilibrium points of the replicator equation (3.9) (Lemma 3.2 in Garay et al. 2018). Furthermore, state (0, 0, 0, 1) at which every individual of the population follows strategy **p**\* is a stable (but not asymptotically stable) rest point of the replicator equation (3.9) (Theorem 4.9 in Varga et al. 2020). If the dynamics (3.9) is restricted to any of the faces with vertex (0, 0, 0, 1) then state (0, 0, 0, 1) is asymptotically stable on that face (Theorem 4.7 in Varga et al. 2020).

One of the eigenvalues at every point of the segment is always 0, the other two of them has zero real part in a single point of the segment which is denoted by **x**_0_ (the upper green point in Figure 2 and 5). Above the point (the part of the segment falling between the states **x**_0_ and (0, 0, 0, 1)) both non-zero eigenvalues have negative real parts while under the point (the part of the segment falling between the states 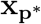 and **x**_0_) the real parts of the two non-zero eigenvalues are positive. To find **x**_0_ we have analyzed the linearization of the right hand side of the dynamical system (3.9) along the segment between 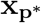 and (0, 0, 0, 1).

We remark that the local property of equilibrium points and the global behaviour of the dynamics on the edges of the phase spaces *S*_3_ and *S*_4_, respectively, are mathematically exact, the eigenvalues were calculated, whereas the global behaviour in the interior of the phase spaces and in the interior of the faces of *S*_4_ are just based on numerical simulations of the phase portrait.

### 3.1 Example 1: Dynamically unstable UESS in rock-scissor-paper game

The popular children’s game, the rock-scissor-paper game describes a game with three pure strategies where each pure strategy beats precisely another pure strategy (rock beats scissor, scissor beats paper and paper beats rock). The payoff matrix has the following form

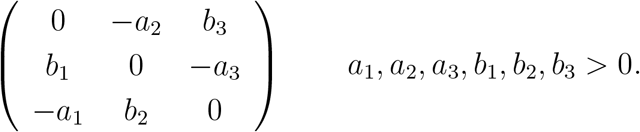

The corresponding replicator equation always has a rest point which is asymptotically stable (and the corresponding mixed strategy is a UESS) if and only if the determinant *b*_1_*b*_2_*b*_3_ − *a*_1_*a*_2_*a*_3_ is positive (Theorem 7.7.2 in Hofbauer and Sigmund 1998). Otherwise, the rest point is a (non-asymptotically) stable center (if the determinant is zero) or unstable (if the determinant is negative). From evolutionary aspect, it models a cyclic dominance (Sinervo and Lively 1996, Kerr et al. 2002, Liao et al. 2019) which can result in the coexistence of the three phenotypes (Hofbauer and Sigmund 1998, Nowak 2006, Mobilia 2010, Toupo and Strogatz 2015).

Now we show that arbitrary small waiting times can destabilize a stable coexistence. Let the payoff matrix *A* = *A*(*s*) and the time constraint matrix *T* = *T* (*s*) be

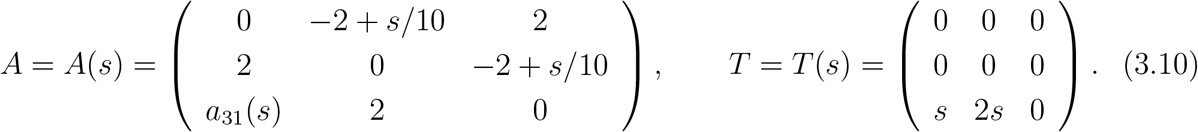

where

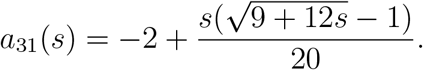

Then for any *s* ∈ (0, *s*_0_) with some *s*_0_ ≥ 3 strategy **p**\* = (1/3, 1/3, 1/3) is a UESS of the matrix game under time constraints with payoff matrix *A*(*s*) and time constraints matrix *T* (*s*) such that the corresponding sate **x**(**p**\*) is an unstable rest point of the replicator equation (3.8). The verification can be found in Appendix A.2.2. Note that the determinant of *A*(*s*) is positive if *s* ∈ (0, 3] so the matrix game with matrix *A*(*s*) (*s* ∈ (0, 3]) has an interior UESS (which tends to **p**\* as *s* → 0) (see expression (7.34) in Hofbauer and Sigmund 1998) and, consequently, **p**\* is an asymptotically stable rest point of the relating replicator equation with respect to pure strategies (Taylor and Yonker 1978, Hofbauer et al. 1979, Zeeman 1980 or Theorem 7.2.4 in Hofbauer and Sigmund 1998).

The phase portrait of dynamics (3.8) for *s* = 1 is depicted in Figure 1. The three vertices together with the edges between them form a heteroclinic cycle. The Jacobian matrix of the right hand side at equilibrium point

**Figure 1.**
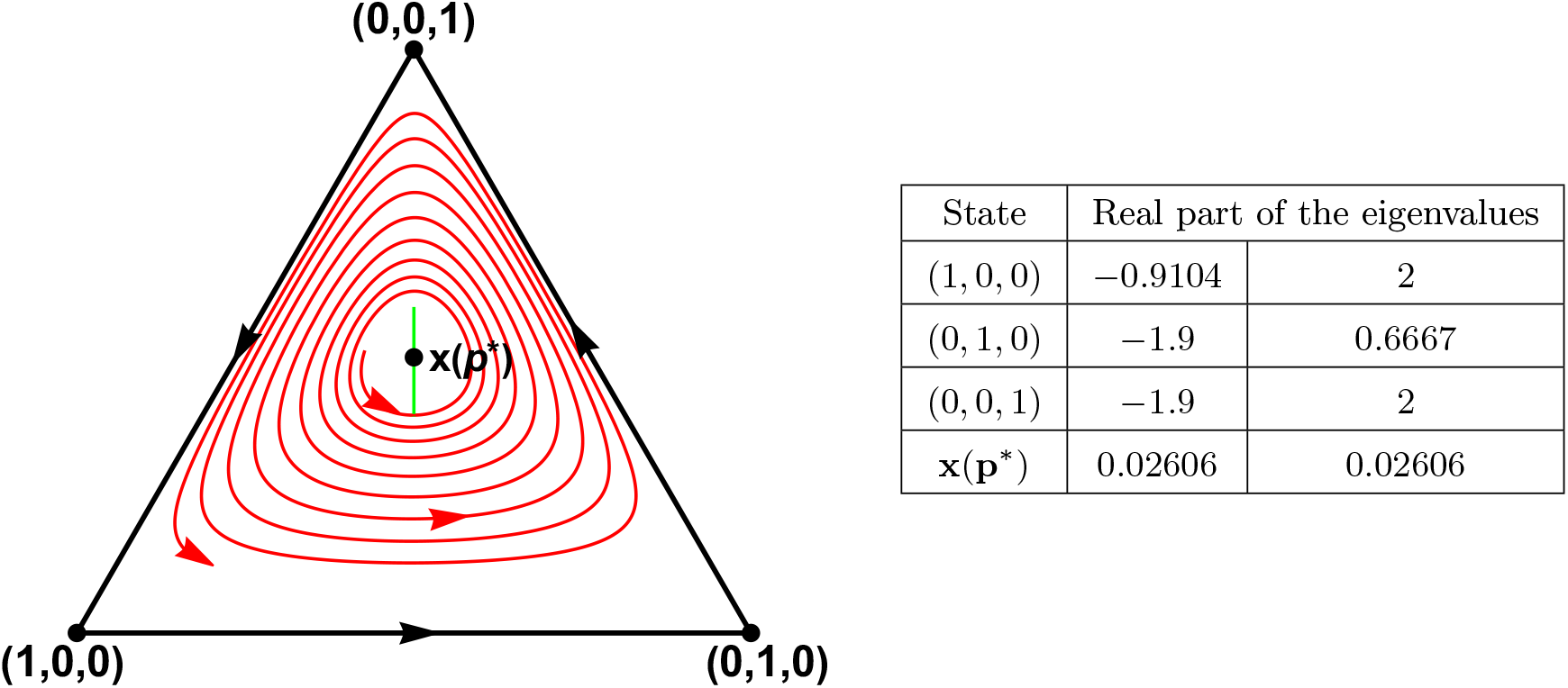
The phase portrait of the replicator equation (3.8) with respect to pure phenotypes **e**_1_, **e**_2_ and **e**_3_. The payoff matrix A and the time constraint matrix T are given in (3.10) setting *s* to be 1. **x**(**p**\*) = **x**^*R*^(**p**\*) is the state corresponding to strategy **p**\* = (1/3, 1/3, 1/3) through Lemma 2.2: 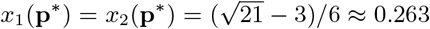. Although **p**\* is a UESS, **x**(**p**\*) is an unstable rest point of the replicator equation. The vertices of the simplex are saddle points and the boundary of the simplex is a heteroclinic cycle which appears to be attractive. For every state **x**, there is a composition 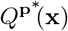 exhibiting strategy **p**\* in the population in state **x** (see section 3.4 for more explanation). Therefore a subpopulation in state 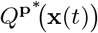 has higher fitness than that of the whole population. The green segment is the set of states 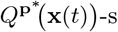 as **x**(*t*) runs over the red orbit (see the enlargement of the green segment in Figure 6). 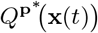 generally differs from **x**(**p**\*). One can intuitively expect that the population “try” to evolve towards 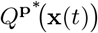 rather than **x**(**p**\*) but 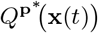 varies from moment to moment which can contribute to the instability of **x**(**p**\*) in the example. In the table, we give the real parts of the eigenvalues of the linearization of the replicator equation at the rest points indicated in the phase portrait.

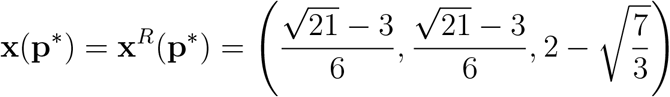

corresponding to the UESS **p**\* = (1/3, 1/3, 1/3) has eigenvalues with positive real part implying the instability of the rest point. The phase portrait suggest that the solutions tend to the boundary of the simplex.

The phase portrait of the extended replicator equation (3.9) is shown in Figure 2. On the face determined by vertices (1, 0, 0, 0), (0, 1, 0, 0), (0, 0, 1, 0), the dynamics agrees with (3.8) so the phase portrait on it is the same as in Figure 1.

**Figure 2.**
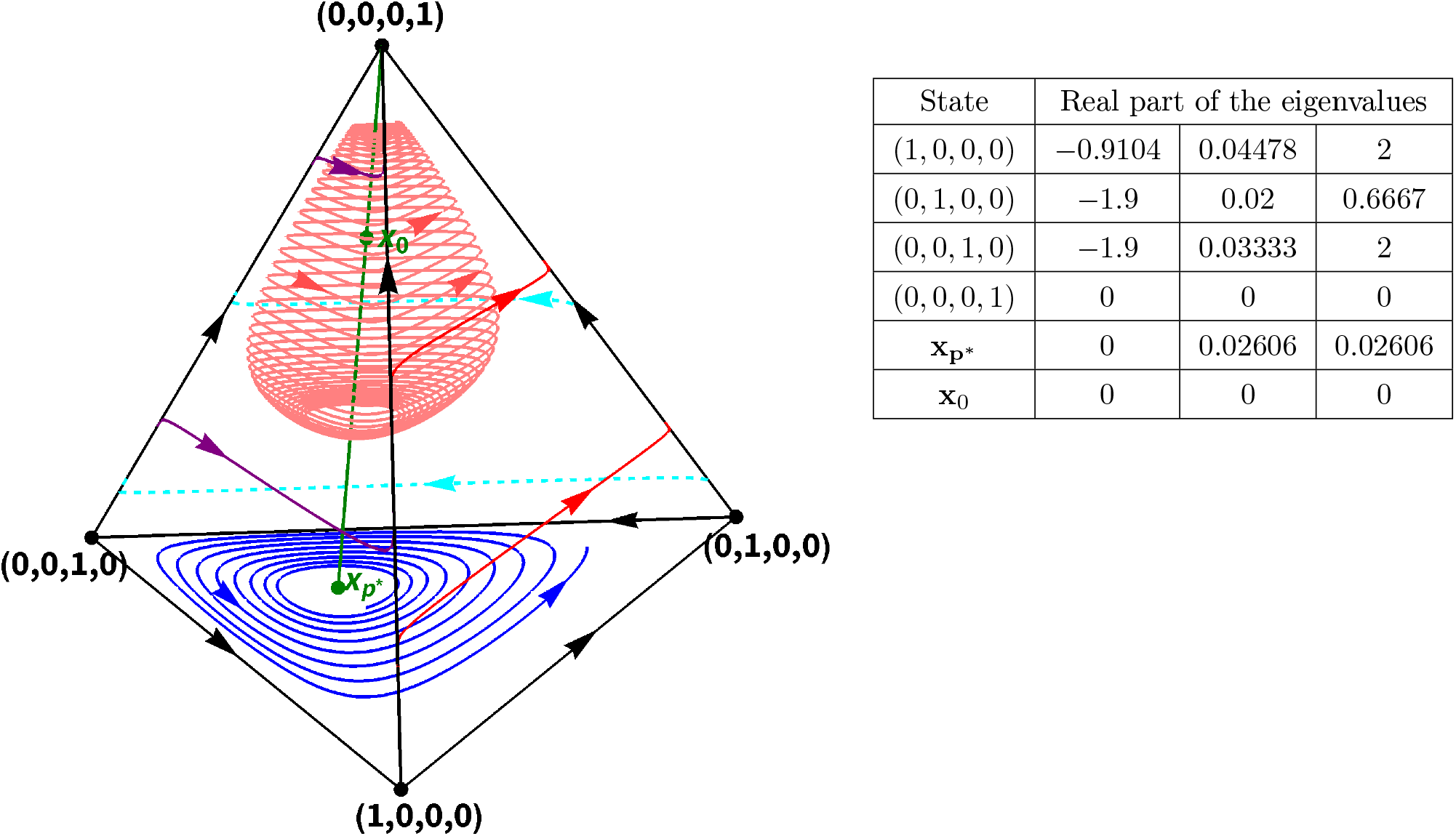
The phase portrait of the replicator equation with respect to phenotypes **e**_1_, **e**_2_, **e**_3_ and **p**\* = (1/3, 1/3, 1/3). The payoff matrix A and the time constraint matrix T are given in (3.10) setting *s* to 1. 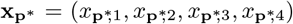 is the state corresponding to strategy **p**\* = (1/3, 1/3, 1/3) through Lemma 2.2 on the face determined by the vertices (1, 0, 0, 0), (0, 1, 0, 0) and (0, 0, 1, 0):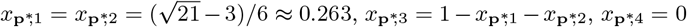. It agrees with **x**(**p**\*) in Figure 1. Every state on the green segment between (0, 0, 0, 1) and 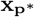 corresponds to strategy **p**\* through Lemma 2.2. Hence every point of the segment is a rest point of the replicator equation. One of the three eigenvalues of the linearization of the replicator equation (3.9) at these states is zero. At **x**_0_, all of the eigenvalues have zero real part. The states on the segment under **x**_0_ have two eigenvalues with positive real part, so they are all unstable though they correspond to the UESS **p**\*. The states between **x**_0_ and (0, 0, 0, 1) whereas have two eigenvalues with negative real part. The state (0, 0, 0, 1) is stable (but not asymptotically). In the table, we give the real parts of the eigenvalues of the linearization of the replicator equation at the rest points indicated in the phase portrait.

The solutions in the interior of the face determined by vertices (1, 0, 0, 0), (0, 1, 0, 0), (0, 0, 0, 1) start from (1, 0, 0, 0) and end in (0, 0, 0, 1). The situation is similar on the other two faces.

The state **x**_0_ on the segment matching 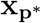 with (0, 0, 0, 1) and corresponding to the UESS **p**\* is equal to **x**^*E*^(*u*_0_/3, *u*_0_/3, *u*_0_/3, 1 −*u*_0_) where 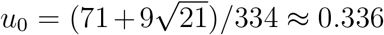. Accordingly, the coordinates of **x**_0_ are (*x*_0_)_1_ ≈ 0.089, (*x*_0_)_2_ ≈ 0.089, (*x*_0_)_3_ ≈ 0.159 and (*x*_0_)_4_ = 1 − *u*_0_ ≈ 0.664.

### 3.2 Limit cycle, Hopf bifurcation

Previously, we have found that both the game with payoff matrix *A*(1) and the game under time constraints with the same payoff matrix and time constraint matrix *T* (1) have a UESS. Nevertheless, the stability properties of the corresponding rest points are different. While the rest point is asymptotically stable in the former case, it is unstable in the latter and the orbits spirally tend to the boundary of the simplex according to the numerical simulation (Figure 1). The situation can remind us of a supercritical Hopf bifurcation. Indeed, let

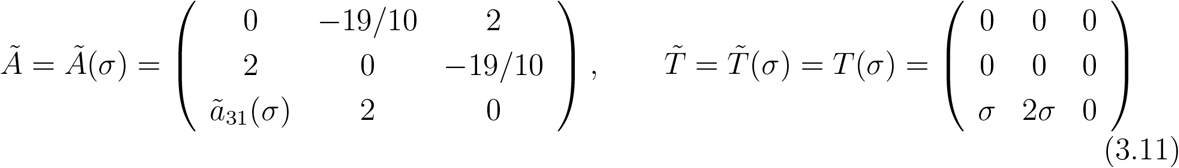

where

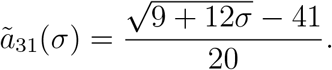

Here *ã*_31_(*σ*) has been chosen such that **p**\* = (1/3, 1/3, 1/3) is a UESS for every *σ* ∈ [0, 1]. Note that *Ã*(1) = *A*(1). Therefore the matrix game under time constraints with matrices *Ã*(1) and 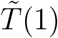 has a UESS such that the corresponding rest point of the replicator equation is unstable while the game with matrices *Ã*(0) and 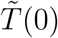 has a UESS such that the corresponding rest point is already asymptotically stable. Further analysis reveals that a supercritical Hopf bifurcation occurs at 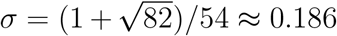 (see Appendix A.3). So a limit cycle around the rest point must exist when *σ* is close to 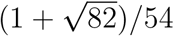. For instance, for *σ* = 19/100, a limit cycle takes shape according to the numerical simulation. However the orbits tending to the limit cycle wind around themselves so tightly that the limit cycle cannot be perceived easily. Therefore, we give a more spectacular example on Figure 3.

**Figure 3.**
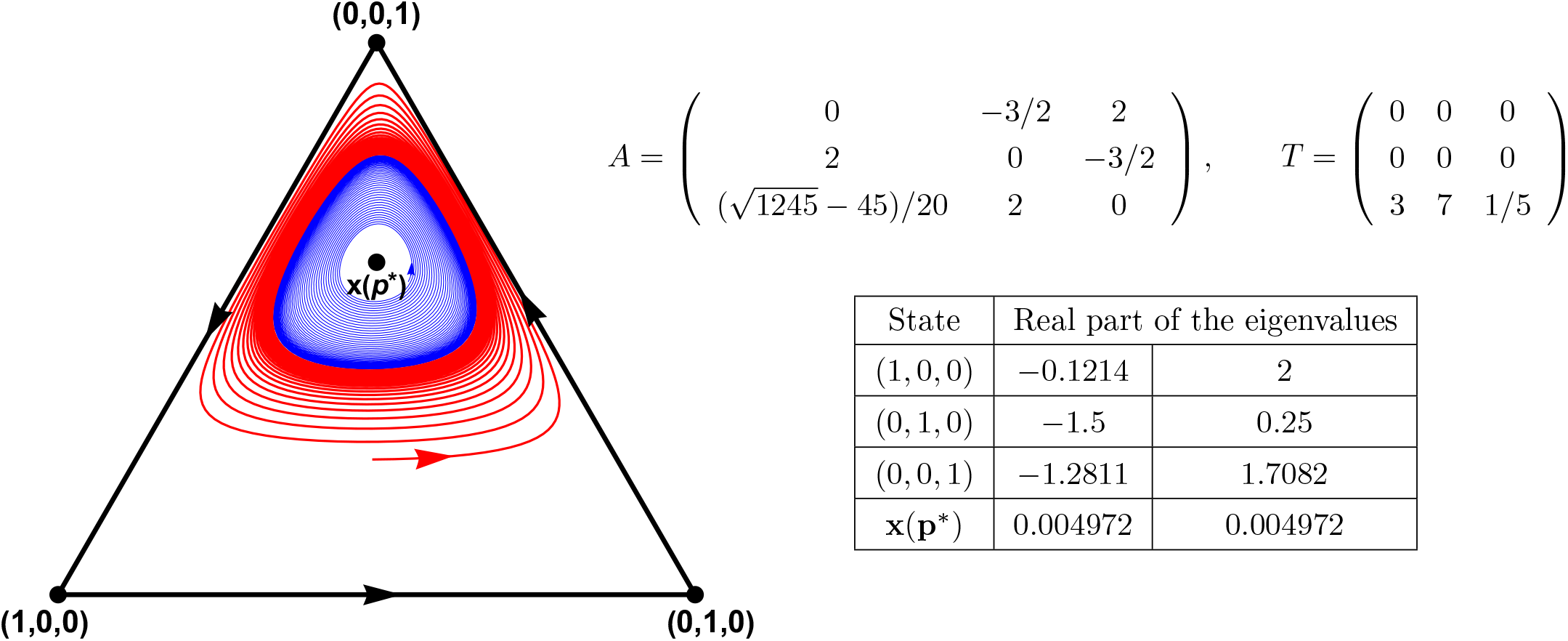
The phase portrait of the replicator equation (3.8) with respect to pure phenotypes **e**_1_, **e**_2_ and **e**_3_. The payoff matrix A and the time constraint matrix T are given right from the phase portrait. **x**(**p**\*) = **x**^*R*^(**p**\*) is the state corresponding to strategy **p**\* = (1/3, 1/3, 1/3) through Lemma 2.2: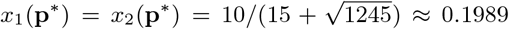. Although **p**\* is a UESS (see Appendix A.2.3), **x**(**p**\*) is an unstable rest point of the replicator equation. The vertices of the simplex are saddle points and the boundary of the simplex is a repelling heteroclinic cycle. It seems that there is a stable limit cycle around **x**(**p**\*). In the table, we give the real parts of the eigenvalues of the linearization of the replicator equation at the rest points indicated in the phase portrait.

### 3.3 Example 2 with a strict NE and a dynamically unstable UESS

The payoff matrix *A* and the time constraint matrix *T* are the following in this case:

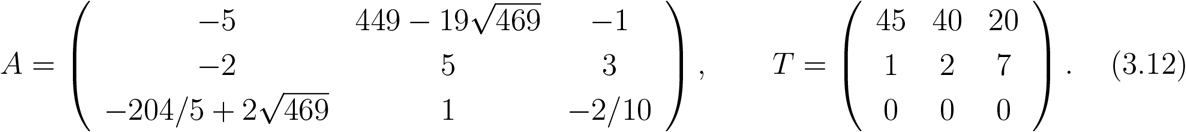

Though similar phase portrait is possible in the classical case (the last element of the first row in Figure 11 in Zeeman 1980, phase portrait 12 in Figure 6 in Bomze 1983), the unstable interior rest point corresponds to a UESS. In addition to **p**\* = (1/3, 1/3, 1/3), there is a further UESS, moreover, a strict Nash equilibrium which is impossible in classic matrix games (see the first paragraph after the proof of Theorem 6.4.1 on p.64 in Hofbauer and Sigmund 1998). A strategy **r** is a strict Nash equilibrium if, for every strategy **q** ≠ **r**, the strict inequality

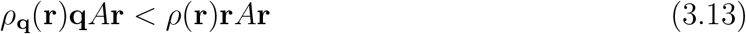

holds (Definition 2.2 in Garay et al. 2018). If **r** is a strict Nash equilibrium then it is a pure strategy (Theorem 4.1 in Garay et al. 2018) and a UESS (Theorem 4.1 in Varga et al. 2020). In the present example, strategy **e**_2_ is a strict Nash equilibrium implying that the corresponding state (0, 1, 0) and (0, 1, 0, 0), respectively, is an asymptotically stable rest point of the replicator equation with respect to **e**_1_, **e**_2_, **e**_3_ and the replicator equation with respect to **e**_1_, **e**_2_, **e**_3_, **p**\*, respectively (Corollary 4.8 in Varga et al. 2020).

To see that **e**_2_ is a strict Nash equilibrium one should check inequality (3.13) with **r** = **e**_2_. By (2.4) and (2.5) we get *ρ*(**e**_2_) and *ρ*_**q**_(**e**_2_), respectively. We obtain that

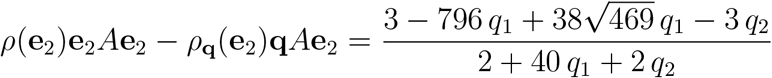

Note that **q** = (*q*_1_, *q*_2_, 1 − *q*_1_ − *q*_2_) ∈ *S*_3_ is a strategy. Thus the denominator 2 + 40*q*_1_ + 2*q*_2_ is positive for every **q**. Consequently, in order to prove that **e**_2_ is a strict Nash equilibrium, it is enough to validate that the numerator 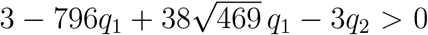 for every **q ≠ e**_2_ which can easily be seen:

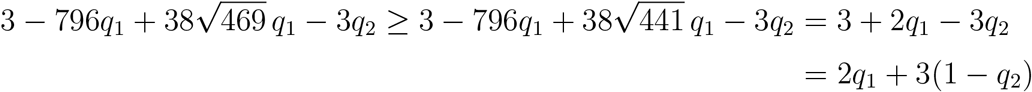

which is positive for every strategy **q** different from **e**_2_. Hence **e**_2_ is a strict Nash equilibrium, as claimed.

Considering the replicator equation with respect to **e**_1_, **e**_2_ and **e**_3_, we find two further rest points besides the vertices and the rest point **x**^*R*^(**p**\*) corresponding to the UESS **p**\*.

There is a rest point on the edge between states (1, 0, 0) and (0, 1, 0). We denote it by **x**^(12)^ ≈ (0.689, 0.311, 0). This is an unstable rest point of the dynamics restricted to the edge and it is a saddle point of the dynamics with respect to the pure strategies.

The other rest point can be found on the edge between (1, 0, 0) and (0, 0, 1). It is denoted by **x**^(13)^ ≈ (0.395, 0, 0.605). This state is asymptotically stable on the edge but it is a saddle point of the replicator dynamics with respect to **e**_1_, **e**_2_ and **e**_3_. A separatrix starts from it and ends in the state (0, 1, 0).

The (interior) solutions on the (1, 0, 0) side of the separatrix start from state **x**(**p**\*) = **x**^*R*^(**p**\*) and end in (0, 1, 0) except for one which ends in **x**^(12)^. The (interior) solutions on the (0, 0, 1) side of the separatrix start from (0, 0, 1) and end in (0, 1, 0) (see Figure 4).

**Figure 4.**
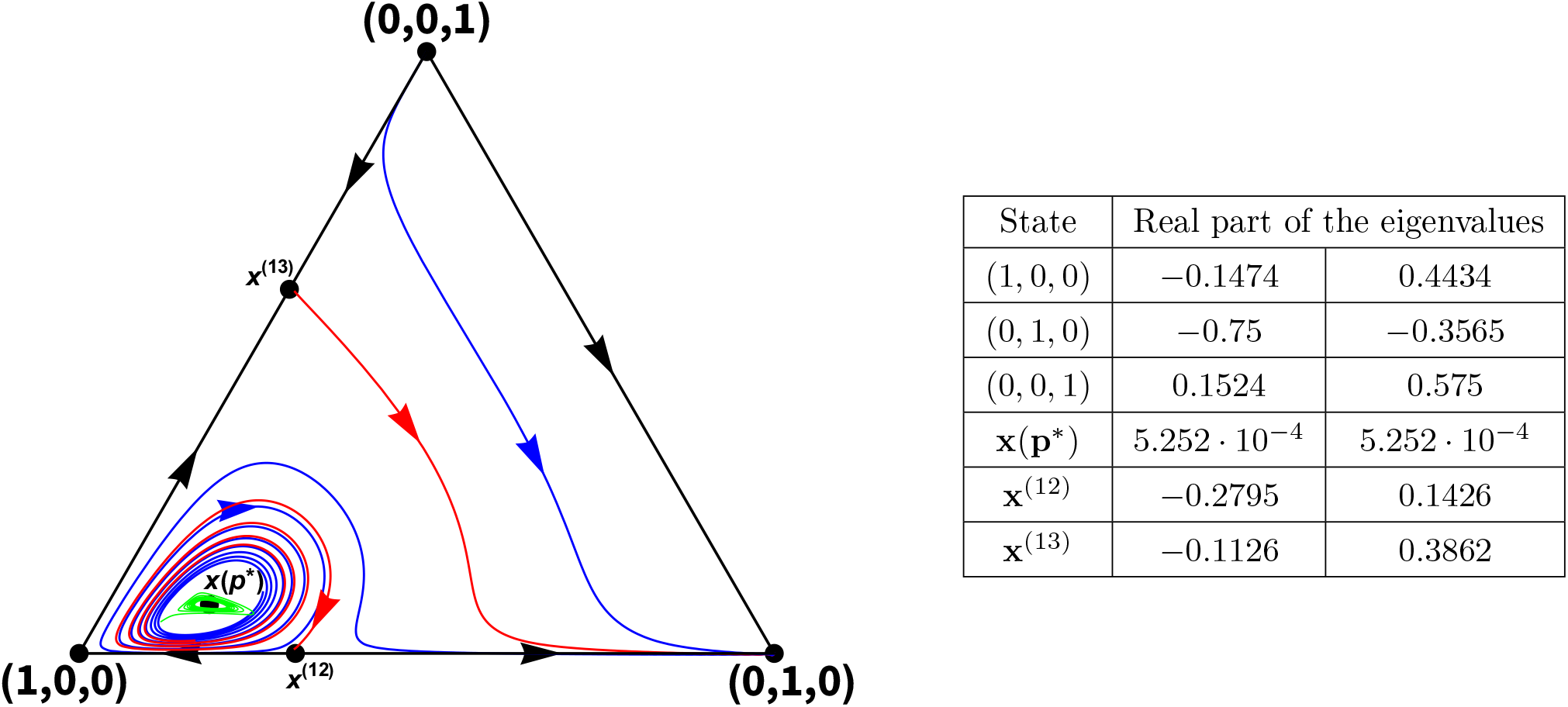
The phase portrait of the replicator equation (3.8) with respect to pure phenotypes **e**_1_, **e**_2_ and **e**_3_. The payoff matrix A and the time constraint matrix T are the matrices in (3.12). **x**(**p**\*) = **x**^*R*^(**p**\*) is the state corresponding to strategy **p**\* = (1/3, 1/3, 1/3) through Lemma 2.2: 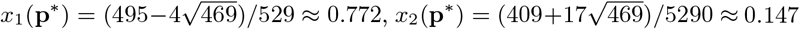. Although **p**\* is a UESS, **x**(**p**\*) is an unstable rest point of the replicator equation. Among the vertices, only (1, 0, 0) is a saddle point; (0, 1, 0) is asymptotically stable, moreover, it corresponds to a strict Nash equilibrium while (0, 0, 1) is a source. In addition to the vertices, **x**^(12)^ and **x**^(13)^ are two further rest points on the boundary of the simplex. Both of them are saddle points. For every state **x**, there is a composition 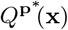 exhibiting strategy **p**\* in the population in state **x** (see section 3.4 for more explanation). Therefore a subpopulation in state 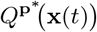 has higher fitness than that of the whole population. The green segment is the set of states 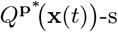 as **x**(*t*) runs over the blue orbit (see the enlargement of the green segment in Figure 6). 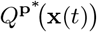 generally differs from **x**(**p**\*). One can intuitively expect that the population “try” to evolve towards 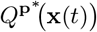 rather than **x**(**p**\*) but 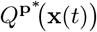 varies from moment to moment which can contribute to the instability of **x**(**p**\*) in the example. In the table, we give the real parts of the eigenvalues of the linearization of the replicator equation at the rest points indicated in the phase portrait.

The coordinates of **x**(**p**\*) are

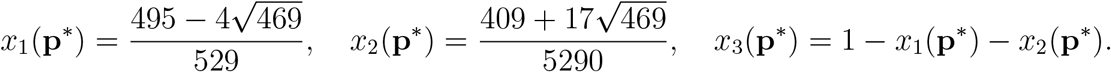

The phase portrait of the extended replicator equation (3.9) is illustrated in Figure 5. The restriction of the dynamics onto the face determined by vertices (1, 0, 0, 0), (0, 1, 0, 0) and (0, 0, 1, 0) is just the replicator equation with respect to pure strategies so the phase portrait on that face agrees with that in Figure 4.

**Figure 5.**
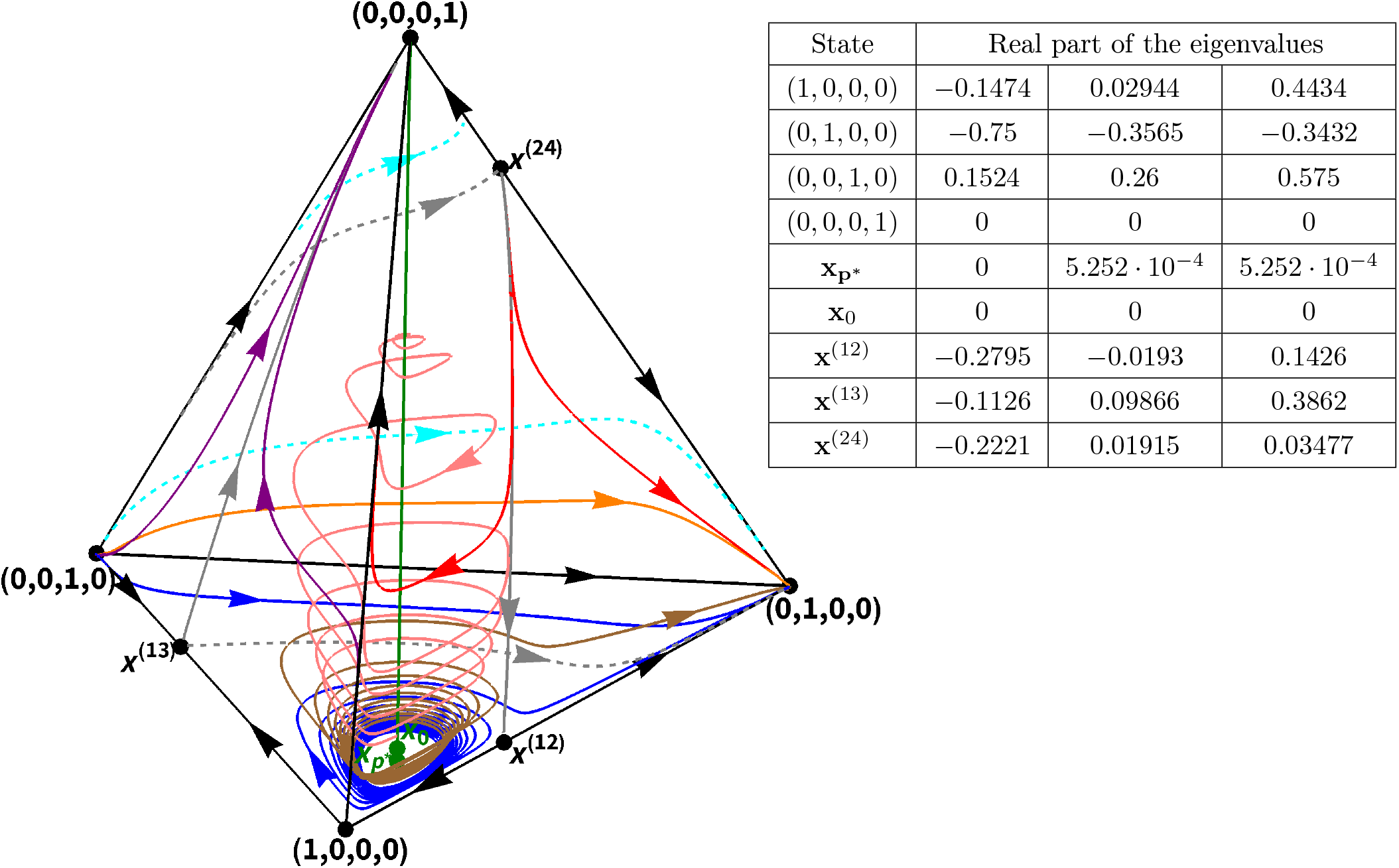
The phase portrait of the replicator equation with respect to phenotypes **e**_1_, **e**_2_, **e**_3_ and **p**\* = (1/3, 1/3, 1/3). The payoff matrix A and the time constraint matrix T are given in (3.12). 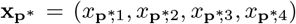 is the state corresponding to strategy **p**\* = (1/3, 1/3, 1/3) through Lemma 2.2 on the face determined by the vertices (1, 0, 0, 0), (0, 1, 0, 0) and (0, 0, 1, 0):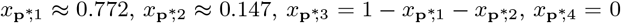. It agrees with **x**(**p**\*) in Figure 4. Every state on the green segment between (0, 0, 0, 1) and 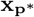 corresponds to strategy **p**\* through Lemma 2.2. Hence every point of the segment is a rest point of the replicator equation. One of the three eigenvalues of the linearization of the replicator equation (3.9) at these states is zero. At **x**_0_, all of the eigenvalues has zero real part. The states on the segment under **x**_0_ have two eigenvalues with positive real part, so they are all unstable though they correspond to the UESS **p**\*. The states between **x**_0_ and (0, 0, 0, 1) whereas have two eigenvalues with negative real part. The state (0, 0, 0, 1) is stable (but not asymptotically). In the table, we give the real parts of the eigenvalues of the linearization of the replicator equation at the rest points indicated in the phase portrait.

There exists a rest point on the edge between (0, 1, 0, 0) and (0, 0, 0, 1). It is denoted by **x**^(24)^. Its coordinates are 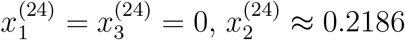, and 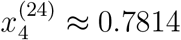. The rest point is unstable with respect to the edge.

It appears that there is a separatrix connecting rest point **x**^(24)^ with rest point **x**^(12)^ on the face determined by the vertices (1, 0, 0, 0), (0, 1, 0, 0) and (0, 0, 0, 1). The separatrix splits the face into two parts. The (interior) orbits start from **x**^(24)^ and end in the stable (moreover asymptotically stable with respect to the face) rest point (0, 0, 0, 1) on the part falling toward the edge between (1, 0, 0, 0) and (0, 0, 0, 1) while in the asymptotically stable rest point (0, 1, 0, 0) on the other part of the face.

Also, it seems that a separatrix runs from **x**^(13)^ to (0, 0, 0, 1) on the face determined by states (1, 0, 0, 0), (0, 0, 1, 0) and (0, 0, 0, 1). The orbits in the interior of the face all end in state (0, 0, 0, 1) but the orbits on the (1, 0, 0, 0) side of the separatrix start from state (1, 0, 0, 0) whereas those falling on the (0, 0, 1, 0) side of the separatrix start from state (0, 0, 1, 0).

It seems that there is also a separatrix on the face determined vertices (0, 1, 0, 0), (0, 0, 1, 0) and (0, 0, 0, 1) connecting state (0, 0, 1, 0) to **x**^(24)^. The orbits in the interior of the face start from (0, 0, 1, 0). The orbits located on the (0, 1, 0, 0) side of the separatrix run into state (0, 1, 0, 0) while those on the (0, 0, 0, 1) side of the separatrix end in sate (0, 0, 0, 1).

In this case, state **x**_0_ is the sate **x**^*E*^(*u*_0_/3, *u*_0_/3, *u*_0_/3, 1 − *u*_0_) where

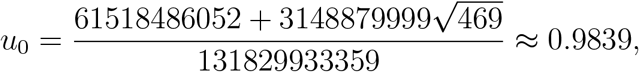

so its coordinates are (*x*_0_)_1_ ≈ 0.7596, (*x*_0_)_2_ ≈ 0.1446, (*x*_0_)_3_ ≈ 0.07981 and (*x*_0_)_4_ = 1 − *u*_0_ ≈ 0.01607.

### 3.4 Composition corresponding to a strategy q in a population of state x

In the frame of classic matrix games, if **p**\* is a UEES then state **p**\* is an asymptotically stable rest point of the replicator equation with respect to pure strategies. The intuitive explanation says that a subpopulation in state **p**\* has a higher mean fitness than the total population of state **x** and therefore the population evolves to state **p**\*. In the morning of classical evolutionary matrix games (Maynard Smith and Price 1973), however the implication was not clear, some years lasted before the discovery of the relation (Taylor and Yonker 1978, Hofbauer et al. 1979, Zeeman 1980).

Our counterexamples show that this classical relation is not self-evident. To help the intuitive comprehension why the implication does not hold in general in the case of matrix games under time constraints, it can be worth to analyse the composition corresponding to the UESS in a population of state **x**.

Lemma 2.2 gives the state **x** that corresponds to a strategy **q** in the sense that the mean strategy of active individuals is just **q** and the proportion of active individuals equals to the proportion of active individuals in the monomorphic population of **q** individuals, in short, **h**(**x**) = **q** and 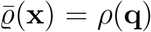. Motivating by this lemma, take a population of pure strategists in state **x** and consider the following composition. If **q** = *q*_1_**e**_1_ + *q*_2_**e**_2_ + *q*_3_**e**_3_ is a strategy then let

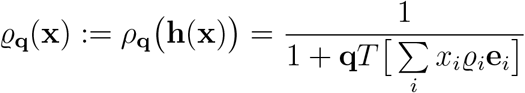

and

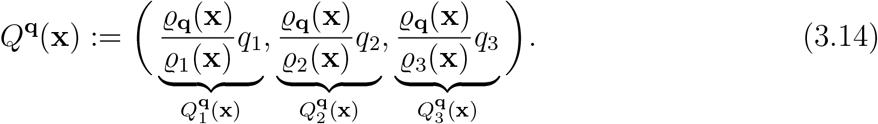

Observe that

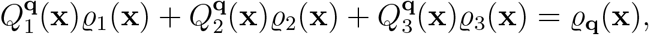

furthermore,

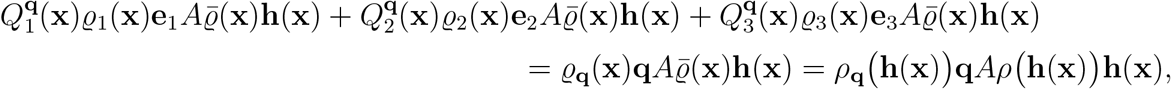

that is, a subpopulation in state *Q*^**q**^(**x**) corresponds to strategy **q** in the sense that the proportion of active individuals in the subpopulation is the same as the probability that a **q** individual playing against the population is active and that the mean fitness of the subpopulation is the same as the fitness of a **q** individual playing against the population.

The observation shows that *Q*^**q**^(**x**) is not a fixed composition in general. It varies with **x**. In particular, if **q** = **p**\*, that is, **q** is a UESS then the subpopulation corresponding to the UESS varies as the state of the population varies (see green curves in Figure 1, 4, 6 and 7). This is a distinction from the calssical case in which the corresponding composition was just the same as the strategy [*Q*^**q**^(**x**) = **q**] independently of **x**. This means that composition 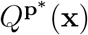 generally differs from **x**(**p**\*) [in the classical case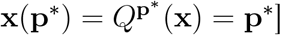. It seems this slight deviation can be enough to make **x**(**p**\*) unstable as shown by our examples.

**Figure 6.**
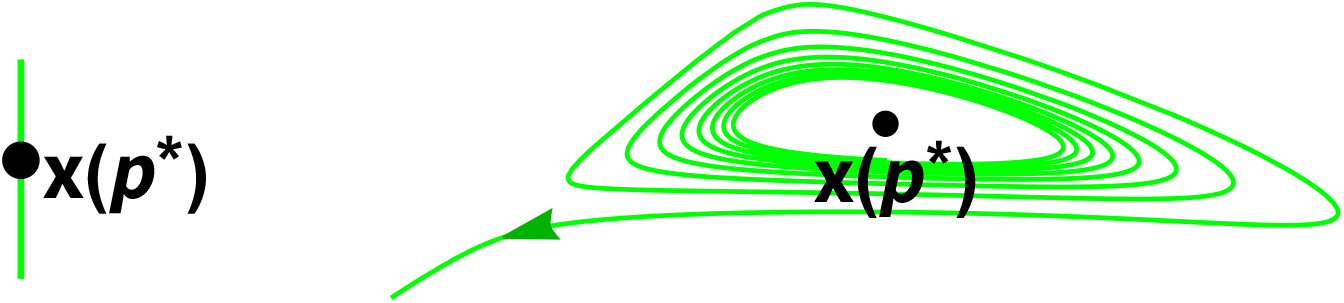
The neighbourhood of **x**(**p**\*) in Figure 1 and 4 are enlarged. The left figure corresponds to Example 1 while the right one to Example 2. The green curves are the states corresponding to **p**\* as **x**(*t*) runs on the orbit around **x**(**p**\*) in Figure 1, 4 (red orbit in Fig. 1, blue orbit in Fig. 2), in other words, states 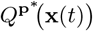 as defined by (3.14) at **q** = **p**\*. If the composition of a subpopulation is 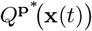 then the mean strategy of active individuals of the subpopulation is just **p**\*. In classic matrix games 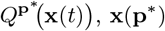 and **p**\* coincide, but not in the examples. This can contribute to the instability of **x**(**p**\*).

**Figure 7.**
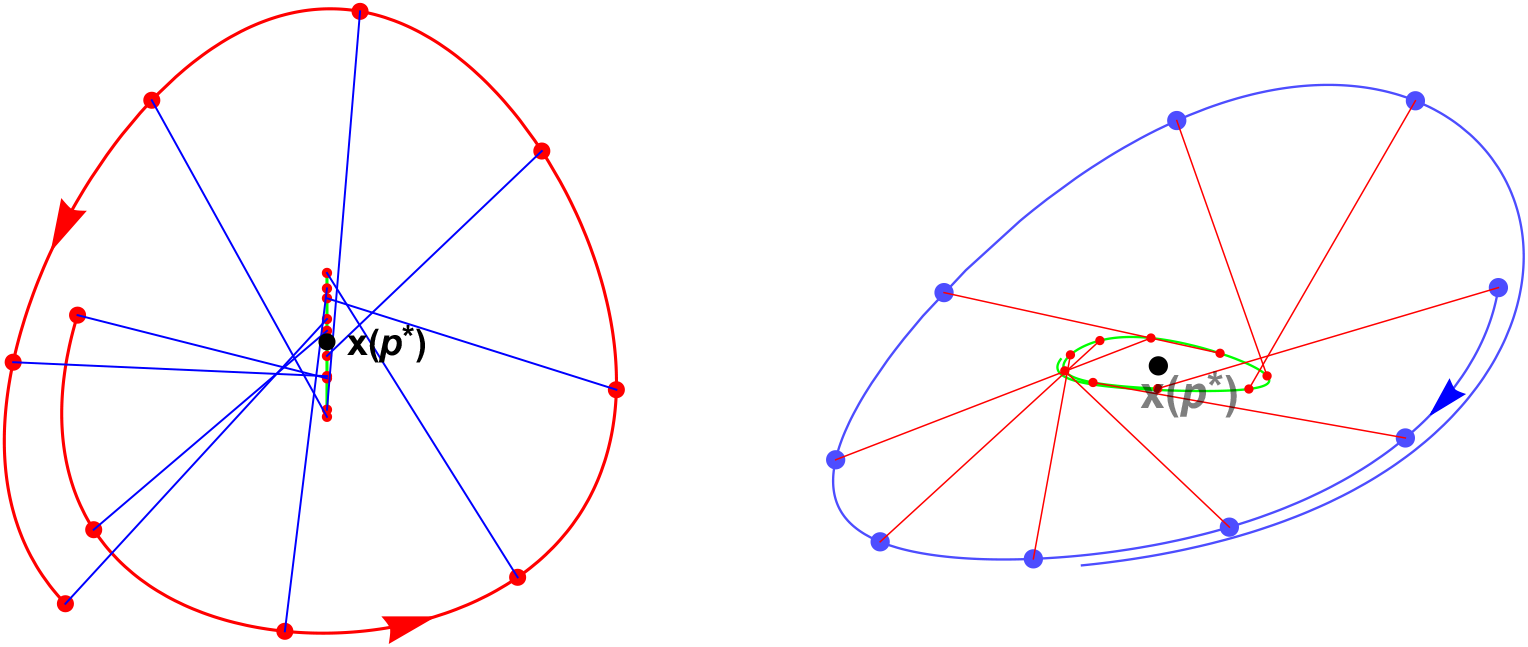
The initial subarc of the orbit **x**(*t*) around **x**(**p**\*) in Figure 1, 4 (red orbit in Fig. 1, blue orbit in Fig. 2) and the corresponding curve drawn by composition 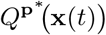 (green curve). If the composition of a subpopulation is 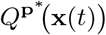 in the population of state **x**(*t*) then the mean strategy of active individuals in the subpopulation is **p**\*. Therefore, one can intuitively expect that the population evolves towards 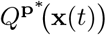 varying from moment to moment rather than **x**(**p**\*) which leads to the instability of **x**(**p**\*) in the examples.

## 4 Discussion

At first glance, analysing the relationship between models on monomorphic and polymorphic populations can primarily appear to be a mathematical problem. However, it is connected with the question of diversity in biology. In the theory of classic evolutionary matrix games, for instance, the existence of an interior monomorphic UESS implies stable diversity in polymorphic situations through strong stability (Cressman 1990, Cressman 1992). Therefore, to make clear the difference between the monomorphic and polymorphic models is an interesting question from the viewpoint of biology, as well.

Garay et al. (2017) and Křivan and Cressman (2017) incorporated time constraints in the model of evolutionary matrix games. As they pointed out, time constraints can essentially change evolutionary outcomes. In this article, we have continued the mathematical analyses started in Garay et al. (2018), Varga et al. (2020) on matrix games under time constraints, and we have demonstrated by two examples that static evolutionary stability does not imply the asymptotic stability of the corresponding state of the replicator equation in three or higher dimensions. This is an essential distinction from the case of classic matrix games where static evolutionary stability implies dynamic stability (Taylor and Yonker 1978, Hofbauer et al. 1979, Zeeman 1980).

Our first example is related to the rock-scissor-paper game. This game has biological relevance in ecology (Sinervo and Lively 1996), microbiology (Kerr et al. 2002) or biotechnology (Liao et al. 2019). Also, it is used for modelling interactions between cancer cells (Tomlinson 1997). In addition to proving that arbitrary small distinctions between waiting time can destabilize the rest point corresponding to a UESS, we pointed out that a stable limit cycle can arise around the rest point. Such a phenomenon is impossible in classic matrix games (Zeeman 1980, Bomze 1983). Mobilia (2010) and Toupo and Strogatz (2015) obtained a similar result but they investigated the effect of mutations rather than time constraints.

The second example exhibited a strict Nash equilibrium in addition to the dynamically unstable interior UESS. This is not possible in the classical case either since the existence of an interior ESS precludes the existence of another Nash equilibrium (see the paragraph after the proof of Theorem 6.4.1 in Hofbauer and Sigmund 1998).

In classic matrix games, if **p**\* is an ESS then state **x**\* with 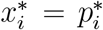 is an evolutionar-ily stable state (Theorem 6.4.1 in Hofbauer and Sigmund 1998) in the population of pure phenotypes, that is, for some *δ* > 0

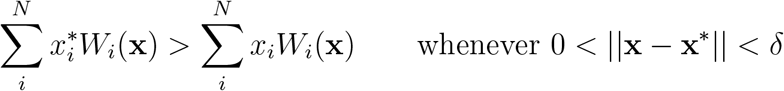

where *W*_*i*_ is the fitness of the *i*-th pure strategy. This condition means that state **x**\* has higher fitness in a slightly perturbed state than the mean fitness of the perturbed state. Therefore, one can expect that the population is steered back to state **x**\*. Indeed, the above relation implies the asymptotic stability of **x**\* with respect to the replicator equation (Hofbauer et al. 1979, Zeeman 1980, Theorem 7.2.4 in Hofbauer and Sigmund 1998). Motivating by this, we investigated the composition corresponding to the UESS in a given population of pure phenotypes. Opposite to the classical case, this composition depends on the current state of the population (Figures 6 and 7). Generally, it does not coincide with the state corresponding to the UESS. This phenomenon can contribute to the instability of the state corresponding to the UESS, as was demonstrated in our examples. On the other hand, this provides only a partial explanation since it is easy to give examples such that the composition corresponding to the UESS varies with the current state of the population, yet the state corresponding to the UESS is asymptotically stable. Therefore future investigations are necessary for a deeper understanding of the stability property of the state corresponding to the UESS.

## A Appendix

### A.1 How to find dynamically unstable UESS

One of the main difficulty in the analyses of the replicator equation (3.7) is that the right hand side generally cannot be given in explicit form. This is because functions *ϱ*_*i*_ defined as the solution of equation system (2.2) cannot be expressed explicitly in general and this problem already araises in three dimension. We first tried examples in which the explicit calculation was possible but this way proved ineffective. Therefore we had to find a more systematic procedure to look for examples with UESS such that the corresponding rest point of the replicator equation is unstable. Here we describe the method used by us. We first quote the following characterization of UESS (Corollary 3.2 in Varga et al. 2020):

*A strategy* **p**\* *is a UESS if and only if there is a δ* > 0 *such that*

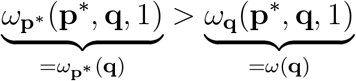

*whenever* 0 < ||**p**\* − **q**|| < *δ*.

Accordingly, if

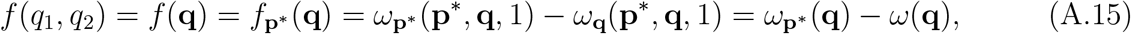

then **p**\* is a UESS if and only if it is a strict local minimum of *f*. Note that since *f* is defined on *S*_3_ which is a 2 dimensional manifold in R^3^, it can be considered as a two variable function of *q*_1_, *q*_2_. It is well-known from multivariable calculus that if

[C1] 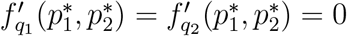,

[C2] 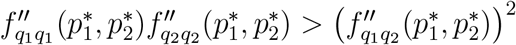 and

[C3] 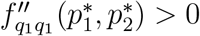 then 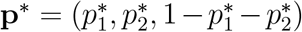 is a strict local minimum of *f*. (see e.g. Theorem 11 of Chapter 14, p. 838 in Thomas 2014). Consequently, if [C1], [C2], [C3] hold for **p**\* = (1/3, 1/3, 1/3) with respect to the matrices *A* and *T*, furthermore, the state **x**(**p**\*) corresponding to **p**\* is an unstable rest point of the replicator equation then we have found a counterexample.

Consider, therefore, the replicator equation (3.8) for the polymorphic population consisting of pure phenotypes **q**_1_ = **e**_1_, **q**_2_ = **e**_2_, **q**_3_ = **e**_3_ with frequencies *x*_1_, *x*_2_ and *x*_3_ = 1 − *x*_1_ − *x*_2_, respectively. Since *x*_1_ + *x*_2_ + *x*_3_ = 1 also holds, any two equations of the system determine the dynamics. It will thus be sufficient to investigate the differential equation system

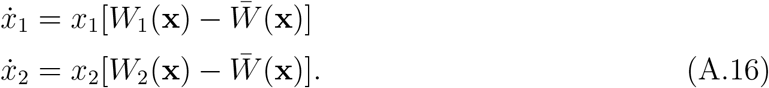

If **p**\* is a NE^3^, then the corresponding state **x**(**p**\*) is a rest point of the replicator equation (Lemma 3.2, Garay et al. 2018). Since (1/3, 1/3, 1/3) is a UESS, which implies that it is a NE too (see the paragraph before Definition 2.2 in Garay et al. 2017), it follows that **x**(1/3, 1/3, 1/3) is a rest point of the replicator equation.

We would like that **x**(1/3, 1/3, 1/3) were unstable. This holds if the Jacobian matrix of the right hand side of (A.16) has a positive eigenvalue at **x**(1/3, 1/3, 1/3). It can be easily checked for a function of two variables that if the characteristic polynomials of the Jacobian matrix is *λ*^2^ + *bλ* + *c* (where *λ* is the variable), then the Jacobian matrix has a positive eigenvalue if and only if

[C4] *b* < 0 or *c* < 0

(see, for example Chapter 4.3 in Kong 2014).

Conditions [C1]-[C4] provide equations and inequalities for the entrances of the payoff matrix *A* and the time constraint matrix *T*. Solving them one can find appropriate matrices *A* and *T* for which there is a UESS **p**\* but the corresponding state **x**(**p**\*) is unstable with respect to the replicator equation.

In connection with [C4] we should avoid the explicit calculation of functions *ϱ*_*i*_ (*i* = 1, 2, 3) because, as mentioned, it is generally not possible. Fortunately, we only need the sign of the coefficients of the characteristic polynomial of the Jacobian matrix at the corresponding state **x**(**p**\*) and this can be calculated. Applying Corollary 2.3, rewrite (A.16) as follows

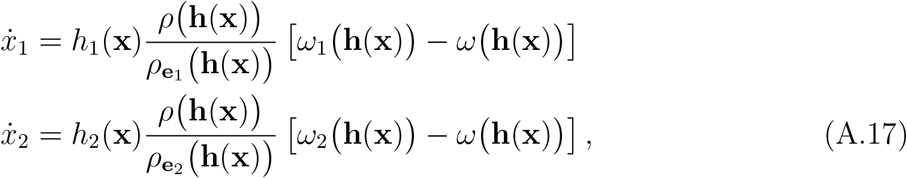

where *h*_1_, *h*_2_ is just the first two components of **h**, that is, **h**(**x**) = *h*_1_ (**x**), *h*_2_ (**x**), 1 − *h*_1_ (**x**) − *h*_2_(**x**)). If

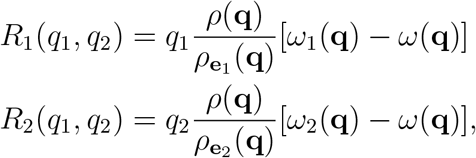

then the Jacobian matrix of the right hand side of (A.17) takes the following shape:

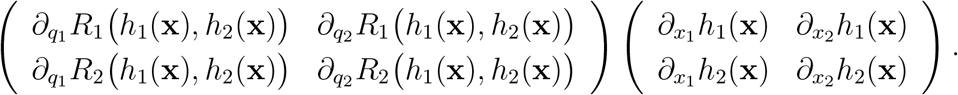

We need the Jacobian at state **x**(1/3, 1/3, 1/3). We therefore express **x** as a function of **q** and then we apply the well-known inverse function theorem (see e.g. Theorem 9.22 and (49) in Chapter 9 (p. 181) in Rudin 1953) to the function **h**(**x**). We obtain that

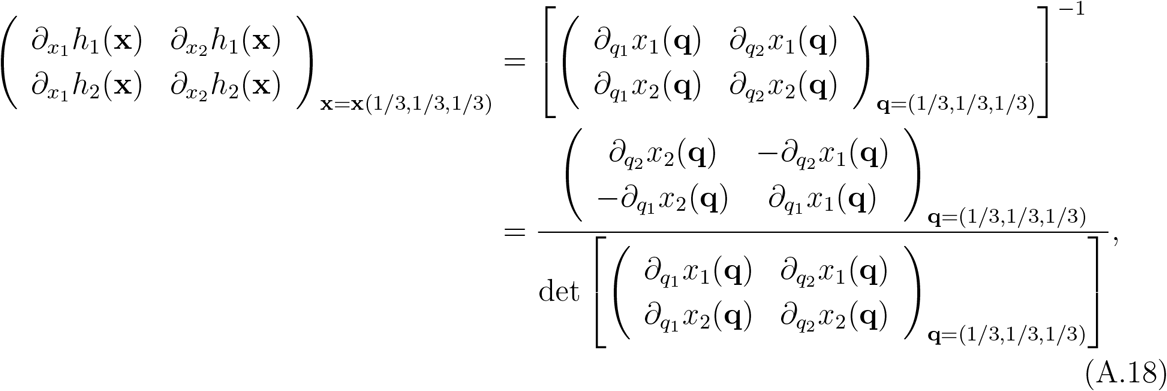

where, of course, the determinant in the denominator of the rightmost side should differ from zero. The rightmost expression already can be calculated and this is enough for our purpose.

As regards dynamics (3.9), one can calculate the linearization of the right hand side similar to that done in (A.18) for replicator equation (3.8). Note that, nevertheless, one should use the maps

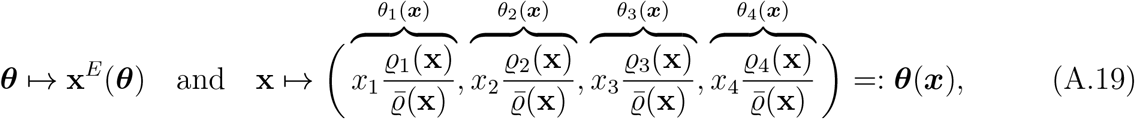

respectively, in this case rather than the maps **p** ↦ **x**^*R*^(**p**) and **x** *1* ↦ **h**(**x**). Accordingly, we rewrite dynamics (3.9) 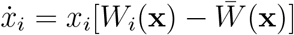 (*i* = 1, 2, 3, 4) as

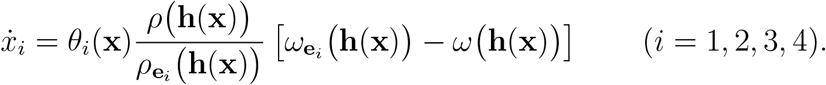

[Note that **h**(**x**) = *θ*_1_(**x**)**e**_1_ + *θ*_2_(**x**)**e**_2_ + *θ*_3_(**x**)**e**_3_ + *θ*_4_(**x**)**p**\* by (2.3) and (A.19).] Also, the functions *R*_*i*_ (*i* = 1, 2, 3) should be given as the functions of the coefficient-list ***θ*** = (*θ*_1_, *θ*_2_, *θ*_3_, *θ*_4_), that is,

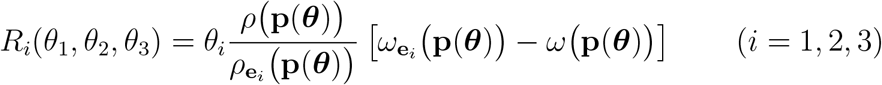

where **p**(***θ***) = **p**(*θ*_1_, *θ*_2_, *θ*_3_) = *θ*_1_**e**_1_ +*θ*_2_**e**_2_ +*θ*_3_**e**_3_ +(1−*θ*_1_ −*θ*_2_ −*θ*_3_)**p**\* (since *θ*_1_ +*θ*_2_ +*θ*_3_ +*θ*_4_ = 1, variable *θ*_4_ can be dropped). Then, similar to (A.18), we can calculate the Jacobian matrix along the segment 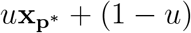 (0, 0, 0, 1), *u* ∈ [0, 1]. After having the linearization along this segment we compute the related characteristic polynomials *λ*^3^ + *b*(*u*)*λ*^2^ + *c*(*u*)*λ* + *d*(*u*). Since one of the eigenvalues is zero at every point of the the segment^4^, it follows that *d*(*u*) = 0 for any *u* ∈ [0, 1]. Therefore the other two eigenvalues should be the zeros of the polynomial *λ*^2^ + *b*(*u*)*λ* + *c*(*u*). The formulas for *b*(*u*) and *c*(*u*), respectively, is not too “beautiful” (we, too, have used Wolfram Mathematica 12), but, after analyzing them, one can see that *b*(0) = *c*(0) = 0 and *c*(*u*) > 0 for *u* ∈ (0, 1]. Hence, the real part of the zeros can be zero only if *b*(*u*) = 0. *b*(*u*) has the form (*b*_11_ −*b*_12_*u*)/(*b*_22_*u*^2^ −*b*_21_*u* +*b*_20_) or *u*(*b*_11_ −*b*_12_*u*)/(*b*_22_*u*^2^ −*b*_21_*u* +*b*_20_) where coefficients *b*_*ij*_ are non-negative, *b*_12_ > *b*_11_ and *b*_20_ > *b*_21_. We infer that the denominator is positive for any *u* ∈ [0, 1] so, if *u* ∈ (0, 1], the sign of *b*(*u*) agrees with the sign of the expression *b*_11_ − *b*_12_*u* which is positive if 0 < *u* < *b*_11_*/b*_12_, negative if *b*_11_*/b*_12_ < *u* ≤ 1 and 0 if *u* = *b*_11_*/b*_12_. It follows (recall condition [C4] too) that the real parts of the two non-zero eigenvalues are positive if 0 < *u* < *b*_11_*/b*_12_, 0 if *u* = *b*_11_*/b*_12_ and negative if *b*_11_*/b*_12_ < *u* ≤ 1. The state belonging to *u* = *b*_11_*/b*_12_ is **x**_0_. The states belonging to *u*-s between 0 and *b*_11_*/b*_12_ forms the segment between 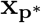 and **x**_0_ while the states belonging to *u*-s between *b*_11_*/b*_12_ and 1 forms the segment between **x**_0_ and (0, 0, 0, 1) (see Figures 2 and 5).

### A.2 Conditions [C1]-[C4] for the examples

In the previous section, we identified some conditions which can be used for seeking examples with dynamically unstable UESS. Here we check the conditions for the examples of the article. Since the expressions arising in the calculation are rather huge we used Mathematica 12 to calculate them and we only give rounded values in the most cases.

#### A.2.1 Conditions [C1]-[C4] for example 1 with *s* = 1

First, we check the special case when *s* = 1. Then one can check that

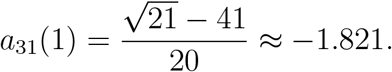

A straightforward calculation yields condition [C1]. For the second order derivatives we obtain:

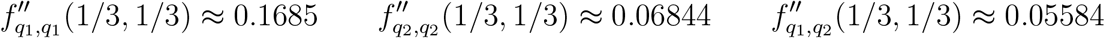

from which we get conditions [C2] and [C3]:

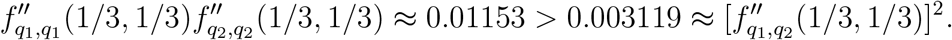

Condition [C4] is true with

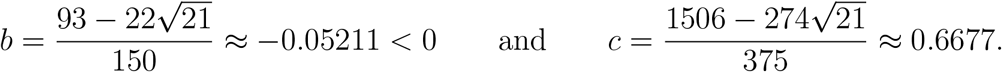

#### A.2.2 Conditions [C1]-[C4] for example 1 with *s* ∈ (0, 3]

Note the following simplification. It is clear that 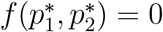 and

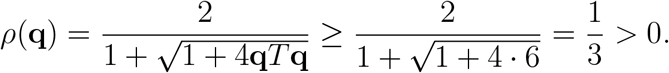

It follows that *f* has a strict minimum at **p**\* if and only if 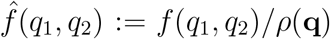 has a strict minimum at **p**\*, so it is enough to check [C1]-[C3] for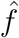. Calculating the first order derivatives we can see that [C1] is valid. For [C2], we get that

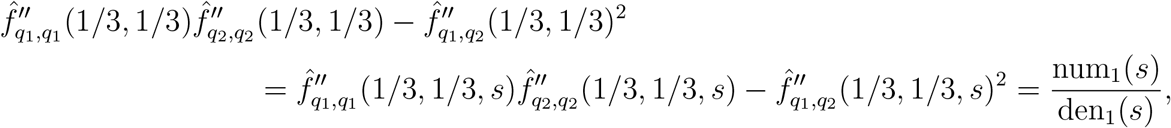

where

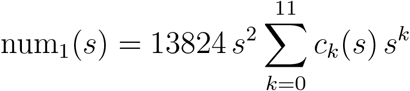

with

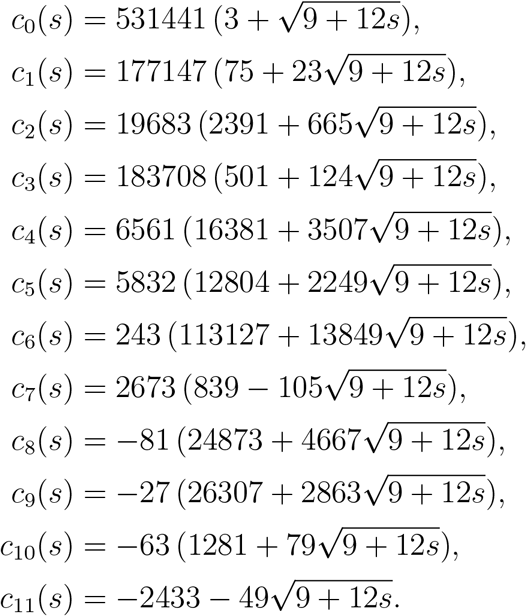

and

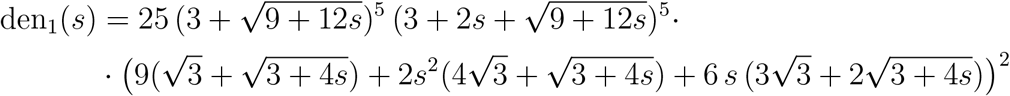

It is easy to see that den_1_(*s*) > 0, if *s* ≥ 0. Also, since *c*_5_(*s*) + *s c*_6_(*s*) > *s*^3^ |*c*_8_(*s*)| + *s*^4^ |*c*_9_(*s*)|, *c*_4_(*s*) > *s*^6^ |*c*_10_(*s*)|, *c*_3_(*s*) > *s*^8^ |*c*_11_(*s*)| and *c*_7_(*s*) > 0 providing that *s* ∈ (0, 3], it immediately follows that num_1_(*s*) > 0 if *s* ∈ (0, 3]. Therefore [C2] holds for *s* ∈ (0, 3].

Furthermore,

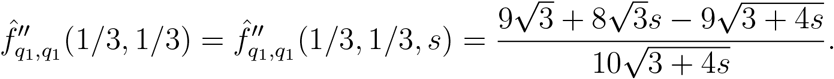

Since

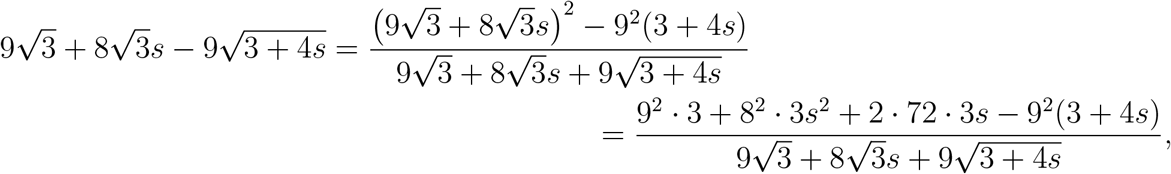

we infer that 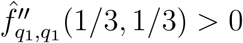 for any *s* > 0, so [C3] holds if *s* ∈ (0, 3].

To check [C4], we simplify the calculation again using the fact that *ρ*(**q**) ≥ 1/3 > 0 and that the orbits of an autonomous differential equation system do not change if each equation is multiplied by the same positive function (Exercise 4.1.3 in Hofbauer and Sigmund 1998).

Therefore we can use differential equation system

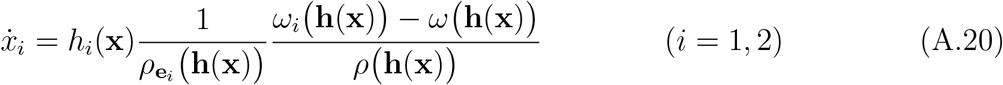

instead of differential equation system (A.17). If the characteristic polynomials of (A.20) at **x**(**p**\*) is 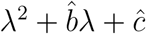 then [C4] is valid if 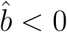 or *ĉ* < 0. We obtain that

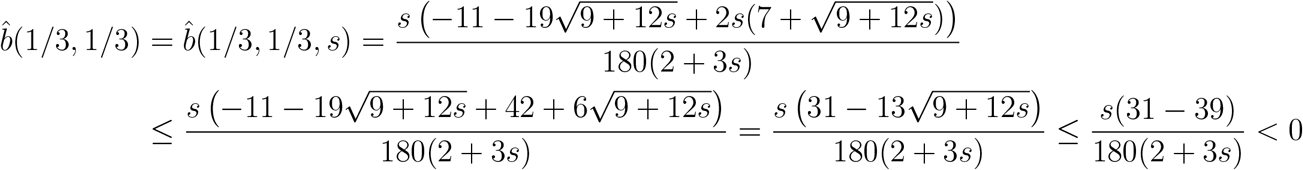

if *s* ∈ (0, 3] and so [C4] is also true.

In summary, [C1]-[C4] hold if *s* ∈ (0, 3] which imply that **p**\* is a UESS such that the corresponding state **x**(**p**\*) is an unstable rest point of the replicator dynamics with respect to pure strategies for any *s* ∈ (0, 3].

#### A.2.3 Conditions [C1]-[C4] for the example in Figure 3

If we calculate the first order derivatives of function *f* in (A.15) we can see that condition [C1] hold. For the second order derivatives, we get

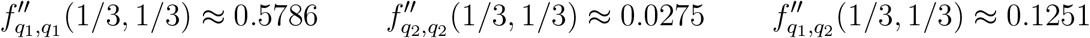

from which we gain that conditions [C2] and [C3] also hold:

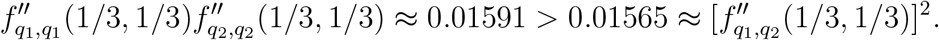

Condition [C4] holds with

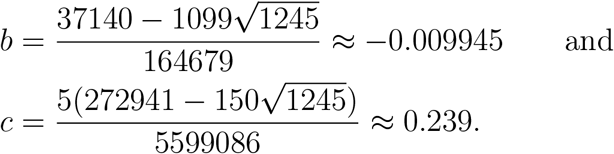

#### A.2.4 Conditions [C1]-[C4] for example 2

Calculating the first order derivatives of function *f* in (A.15) at **q** = (1/3, 1/3) we get that both of them are zero so condition [C1] is valid. The second order derivatives are

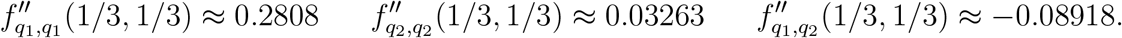

Hence we infer that conditions [C2] and [C3] are also satisfied:

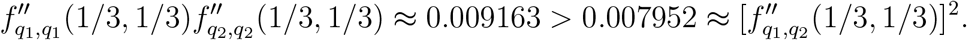

Condition [C4] holds with

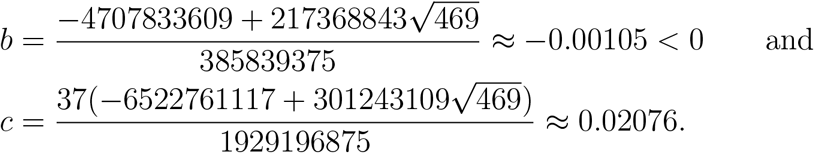

### A.3 Supercritical Hopf bifurcation

Here we check that Hopf bifurcation occurs as *σ* runs from 0 to 1 in the replicator equation associated to the game with payoff matrix *Ã*(*σ*) and time constraint matrix 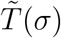 defined in (3.11).

For this purpose, consider a two dimensional system:

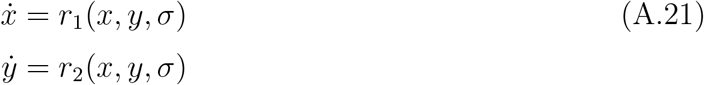

where *σ* ∈ (*σ*_1_, *σ*_2_) is a parameter, *r*_1_ and *r*_2_ are continuously differentiable at least five times. Assume that (*x*\*, *y*\*) is an equilibrium point for every *σ* ∈ (*σ*_1_, *σ*_2_) and there is a *σ*_0_ ∈ (*σ*_1_, *σ*_2_) such that the Jacobian of the right hand side has purely imaginary eigenvalues that is *r*_1_(*x*\*, *y*\*, *σ*) = *r*_2_(*x*\*, *y*\*, *σ*) = 0 and the eigenvalues of

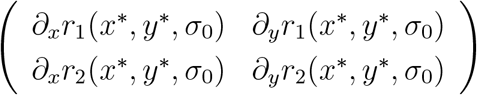

are ±*iβ*(*σ*_0_) with *β*(*σ*_0_) > 0. Denote by *B*(*σ*) the Jacobian

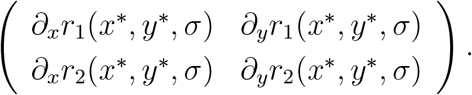

Then the right hand side can be rewritten in the following form:

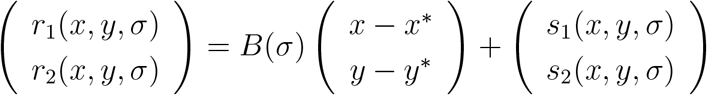

where *s*_*i*_(*x*\*, *y*\*, *σ*) = *∂*_*x*_*s*_*i*_(*x*\*, *y*\*, *σ*) = *∂*_*y*_*s*_*i*_(*x*\*, *y*\*, *σ*) = 0 (*i* = 1, 2).

If *α*(*σ*) ± *iβ*(*σ*) are the eigenvalues of *B*(*σ*) and

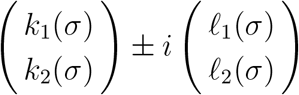

are the corresponding eigenvectors then let

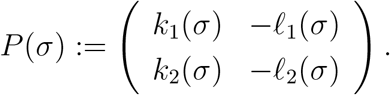

Introduce the new variables *u, v* such that

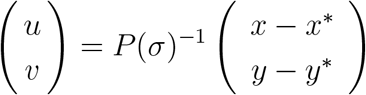

and let

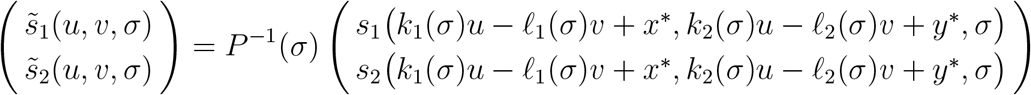

Using variables *u, v*, system (A.21) is transformed into the following form

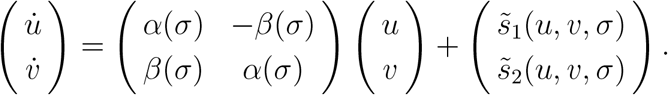

Consider the so-called Lyapunov coefficient as follows:

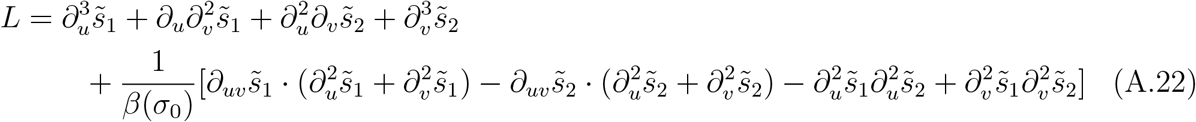

where the partial derivatives are taken at (*u, v, σ*) = (0, 0, *σ*_0_)

The following theorem ensures the occurrence of a supercritical Hopf bifurcation at *σ*_0_.

#### Theorem A.1

(Theorem 3.1.3 in Wiggins (2003) or Theorem 3.4.2 in Guckenheimer (1983)) *Using the above notation, assume that the function α*(*σ*) *is differentiable, β*(*σ*) *is continues, α*(*σ*_0_) = 0, *α*′(*σ*_0_) ≠ 0 *and β*(*σ*_0_) ≠ 0. *If L* < 0 *then a supercritical Andronov-Hopf bifurcation occurs at σ* = *σ*_0_, *that is*,

- *if α*′(*σ*_0_) > 0 *then* (*x*\*, *y*\*) *is asymptotically stable rest point for σ* < *σ*_0_ *while unstable for σ* > *σ*_0_, *with an asymptotically stable periodic orbit around* (*x*\*, *y*\*) *for σ* > *σ*_0_;
- *if α*′(*σ*_0_) < 0 *then* (*x*\*, *y*\*) *is asymptotically stable rest point for σ* > *σ*_0_ *while unstable for σ* < *σ*_0_, *with an asymptotically stable periodic orbit around* (*x*\*, *y*\*) *for σ* < *σ*_0_.

We apply the previous theorem for our example. Again, since expressions arising in calculations are huge, we used Mathematica 12. We start from replicator equation (3.8) associated with payoff matrix *Ã*(*σ*) and time constraint matrix 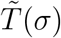 defined in (3.11). Since *x*_3_ = 1 − *x*_1_ − *x*_2_, the third equation can be neglected. If *x* = *x*_1_ and *y* = *x*_2_ then 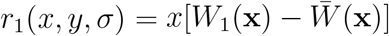 and 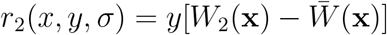. We get that

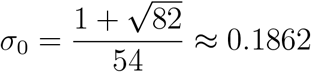

The characteristic polynomial at *σ*_0_ is

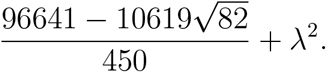

It immediately follows that *α*(*σ*_0_) = 0 and *β*(*σ*_0_) ≠ 0. In general, the characteristic polynomial is *λ*^2^ + *b*(*σ*)*λ* + *c*(*σ*) with

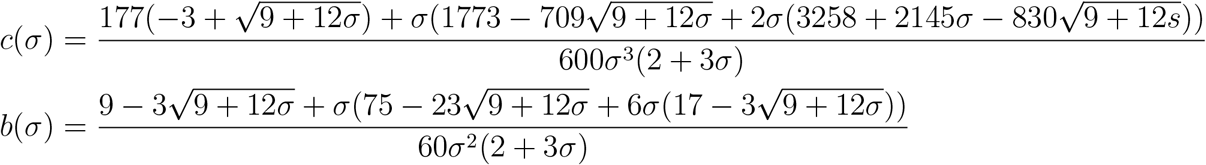

Since

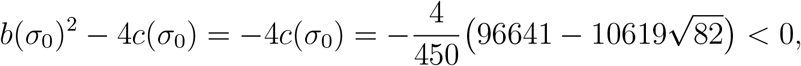

it follows that *b*(*σ*)^2^ − 4*c*(*σ*) < 0 also holds in a neighbourhood of *σ*_0_. For such *σ*-s we have that *α*(*σ*) = −*b*(*σ*)/2 and so we get that

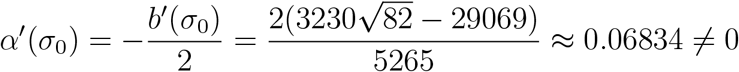

For the Lyapunov coefficient (A.22), we obtain an enormous expression so here we give a rounded value:

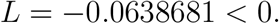

(Note that the precise value is not interesting but the sign which is negative according to Mathematica 12.) In summary, the assumption of Theorem A.1 are satisfied implying that supercritical Hopf bifurcation occurs at *σ*_0_.

## Declarations

### Ethical Approval

Not applicable

### Competing interests

The authors have no conflicts of interest to declare.

## Authors’ contributions

TV and JG designed the study and wrote the main manuscript text, TV constructed and analysed the examples, TV prepared the figures, TV wrote the appendix. Both authors reviewed the manuscript.

## Funding

This research was supported by NKFIH, Hungary KKP 129877 (to TV).

## Availability of data and materials

All data generated or analysed during this study are included in this published article.

## Footnotes

1 The uniqueness trivially follows from the fact that every **r** = (*r*_1_, *r*_2_, *r*_3_) ∈ *S*_3_ is a unique convex combination of the pure strategies: **r** = *r*_1_**e**_1_ + *r*_2_**e**_2_ + *r*_3_**e**_3_.

2 It corresponds to the case when each entrance of *T* is the same, say, 0.

3 A strategy **p**\* is called Nash equilibrium for the matrix game under time constraints (NE in short) if the inequality 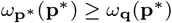 holds for any strategy **q** ∈ *S*_*N*_ (Definition 2.2 in Garay et al. 2018).

4 This is because every point of the segment is a rest point of the dynamics (3.9).

## Notes

### Competing Interest Statement

The authors have declared no competing interest.

### Summary of Updates

Example 1 in Section 3.1 has been replaced with a new example featuring the payoff matrix of a rock-scissors-paper game. Example 2 in Section 3.2 of the previous version has been removed and replaced with an example demonstrating Hopf bifurcation, confirming the possibility of an isolated periodic solution. Example 3 in Section 3.3 of the previous version remains in the manuscript, but it is now renumbered as Example 2. Some calculations and reasoning have been moved to an appendix.

